# An active chromatin interactome in relevant cell lines elucidates biological mechanisms at genetic risk loci for dermatological traits

**DOI:** 10.1101/2020.03.05.973271

**Authors:** Chenfu Shi, Helen Ray-Jones, James Ding, Kate Duffus, Yao Fu, Vasanthi Priyadarshini Gaddi, Oliver Gough, Jenny Hankinson, Paul Martin, Amanda McGovern, Annie Yarwood, Patrick Gaffney, Steve Eyre, Magnus Rattray, Richard B Warren, Gisela Orozco

**Affiliations:** Centre for Genetics and Genomics Versus Arthritis. Division of Musculoskeletal and Dermatological Sciences, School of Biological Sciences, Faculty of Biology, Medicine and Health, The University of Manchester, UK; Dermatology Centre, Salford Royal NHS Foundation Trust, Manchester NIHR Biomedical Research Centre, Manchester Academic Health Science Centre, Manchester, UK; Genes and Human Disease Research Program, Oklahoma Medical Research Foundation, Oklahoma City, 73104, OK, USA; Division of Infection, Immunity and Respiratory Medicine, School of Biological Sciences, University of Manchester, Manchester, UK; The Lydia Becker Institute of Immunology and Inflammation, University of Manchester, Manchester, UK; NIHR Manchester Biomedical Research Centre, Manchester University NHS Foundation Trust, Manchester Academic Health Science Centre, Manchester, UK; Division of Informatics, Imaging and Data Sciences, Faculty of Biology, Medicine and Health, University of Manchester, UK

**Author notes:** These authors contributed equally to this manuscript (Chenfu Shi, Helen Ray-Jones).

## Abstract

Chromatin looping between regulatory elements and gene promoters presents a potential mechanism whereby disease risk variants affect their target genes. Here we use H3K27ac HiChIP, a method for assaying the active chromatin interactome in two cell lines: keratinocytes and skin derived CD8+ T cells. We integrate public datasets for a lymphoblastoid cell line and primary CD4+ T cells and identify gene targets at risk loci for skin-related disorders. Interacting genes enrich for pathways of known importance in each trait, such as cytokine response (psoriatic arthritis, psoriasis) and replicative senescence (melanoma). We show examples of how our analysis can inform changes in the current understanding of multiple psoriasis associated risk loci. For example, the variant rs10794648, which is generally assigned to *IFNLR1*, was linked to *GRHL3* in our dataset, a gene essential in skin repair and development. Our findings, therefore, indicate a renewed importance of skin related factors in the risk of disease.

Graphical Abstract
In this article we take disease associated variants from 5 dermatological conditions and use cell type specific datasets to map genes that could be affected by these variants, providing insight into disease mechanisms.

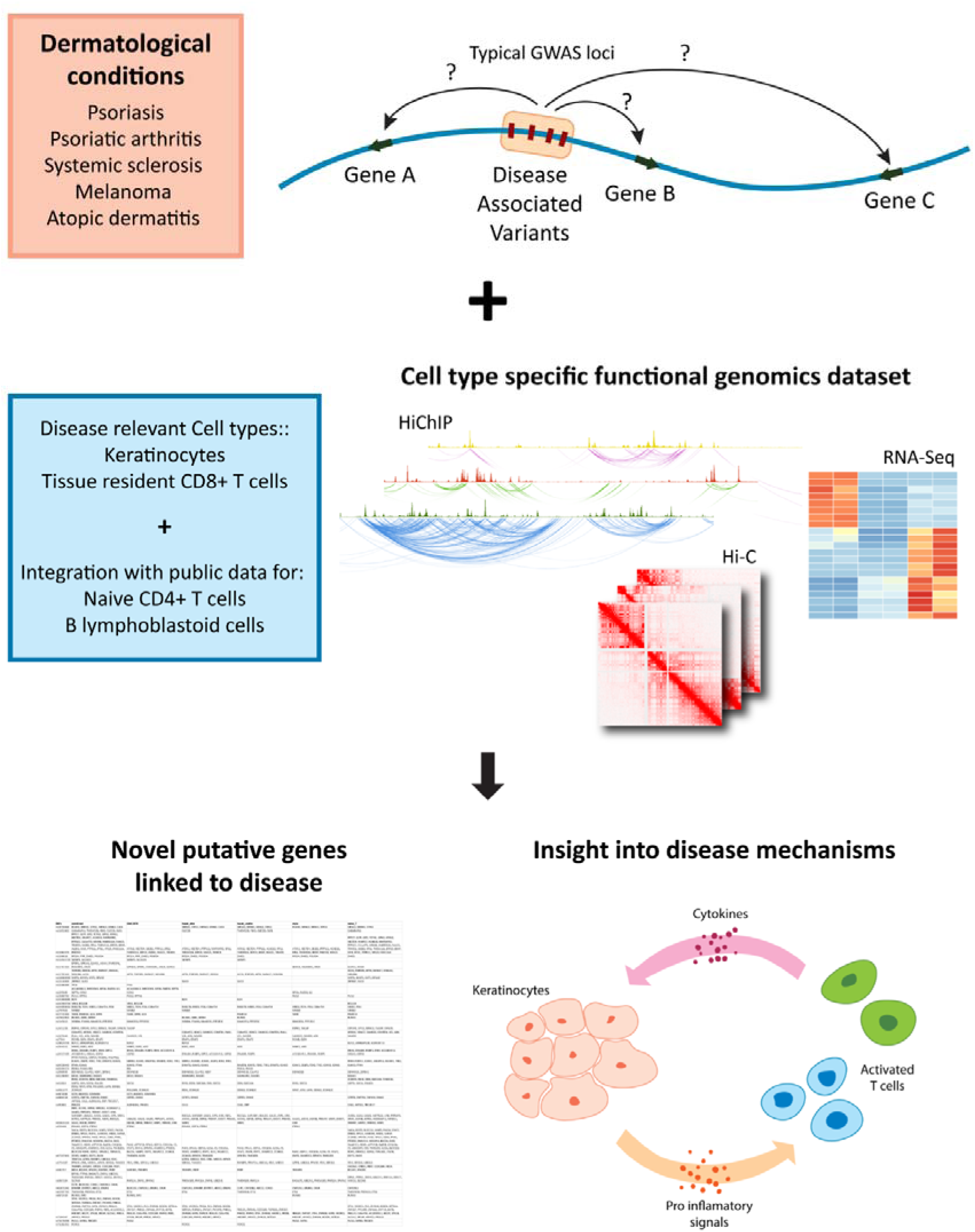

## Introduction

Genome wide association studies (GWAS) have uncovered the genetic factors that contribute to disease risk for many complex disorders. It is now accepted that the majority of these genetic risk factors do not influence coding sequences directly but rather regulatory regions such as enhancers and promoters which can be highly cell type specific (Ernst et al., 2011; Farh et al., 2015; Kundaje et al., 2015). Many studies have also demonstrated that the effect of these variants is not necessarily mediated by the gene that is closest to them as they can affect genes that are very far away from their genomic location via chromatin looping mechanisms (Aguet et al., 2017; GTEx Consortium, 2013; Javierre et al., 2016; Nica & Dermitzakis, 2013; Rao et al., 2014; Võsa et al., 2018).

Recently, there has been a growing interest in using chromatin conformation and other functional genomics techniques to investigate these disease-associated loci and identify the genes that are affected by them. Previous studies have used techniques such as Capture Hi-C and HiChIP to link the genes that physically interact with disease associated loci (Cairns et al., 2016; Dryden et al., 2014; Jäger et al., 2015; P. Martin et al., 2016, 2015; McGovern et al., 2016; Mumbach et al., 2017). These studies have mainly focused on cells derived from blood and immune cells. In the context of dermatological traits, for instance, the H3K27ac-mediated interactome was recently explored in CD4+, T helper (Th) 17 and regulatory T cell subsets (Jeng et al., 2019).

Multiple publications have shown that interactions are cell type specific and are altered during differentiation and stimulation (Burren et al., 2017; Dixon et al., 2012; Hansen, Cattoglio, Darzacq, & Tjian, 2018; Mumbach et al., 2017; Rao et al., 2014; Rubin et al., 2017; Schmitt et al., 2016; Siersbæk et al., 2017). Although for many autoimmune diseases these genetic factors primarily affect immune cells, other cell types might be involved in the disease development (Farh et al., 2015; Mahajan et al., 2018; Mizoguchi et al., 2018). Autoimmune dermatological traits such as psoriasis (Ps), psoriatic arthritis (PsA) and systemic sclerosis (SSc) are systemic conditions with heterogeneous effects in multiple cell types and are likely to involve a complex interplay between skin-resident cells and immune cells (Albanesi, Madonna, Gisondi, & Girolomoni, 2018; Mccoy et al., 2017)

GWAS have revealed that most inherited inflammatory diseases have a shared genetic background, but some conditions are more genetically similar than others. Meanwhile some diseases cannot be easily differentiated based on genetics; for example, Ps and PsA only seem to differ at a small number of loci (Bowes et al., 2017; Stuart et al., 2015). SSc and RA also share a large proportion of genetic risk (López-Isac et al., 2017). Meanwhile, for example, Ps and atopic dermatitis have distinct genetic risk factors and even have opposing genetic mechanisms at some loci (Baurecht et al., 2015). Our knowledge of the differing genetic mechanisms between diseases is hampered by limitations in GWAS sample size, but also in our understanding of how the risk variants influence biological processes within relevant cell types and conditions.

In Ps, the focus in recent years has been on immune cells. There are two T helper cell subsets that are particularly important in disease: Th1 (IFN-γ, IL-12) and Th17 (IL-17A, IL-17F, IL-22) (Lowes et al., 2008). However, recently there has been a rejuvenated interest in the response of keratinocytes, which are the most predominant cells in the epidermis and highly dysregulated in disease in terms of proliferation and differentiation (X. Ni & Lai, 2020). Keratinocytes respond to T cell signals by producing pro-inflammatory cytokines that further contribute to T cell activation (Albanesi et al., 2018; Benhadou, Mintoff, Schnebert, & Thio, 2018; Lorscheid et al., 2019). As a consequence of this, interferon gamma (IFN-γ) stimulated keratinocytes are of interest. IFN-γ is an inflammatory mediator implicated in multiple immune-mediated conditions such as SSc (Wu & Assassi, 2013), Ps and others (Lees, 2015). In Ps in particular, psoriatic lesions are highly enriched in IFN-γ (Schlaak et al., 1994; Uyemura, Yamamura, Fivenson, Modlin, & Nickoloff, 1993). IFN-γ promotes epidermal keratinocyte apoptosis (Hijnen et al., 2013) and represents the classical Th1 pathway whereby IFN-γ is released in abundance by activated T cells in psoriatic skin (Lowes et al., 2008).

For many inflammatory skin diseases, such as melanoma and psoriasis, amongst others, a recurrent feature is the invasion of CD8^+^ T cells in the inflammation site (Antohe et al., 2019; Cai, Fleming, & Yan, 2012; Hennino et al., 2007; O’reilly, Hü Gle, & Van Laar, n.d.). In psoriasis, these cells release IL17, an important factor in disease pathology (Ortega et al., 2009). Tissue resident CD8^+^ T cells can be altered and have different phenotypes compared to circulating/naïve cells. For inflammatory diseases with high skin involvement, studying interactions from Hi-C maps in skin derived CD8+ T cells, as well as resting and IFN-γ stimulated keratinocytes would be valuable in advancing the study of disease mechanisms, but the amount of sequencing required to fully map genome-wide interactions across all cell types is technically challenging.

HiChIP is a recently developed technique that allows detection of long range interactions with increased sensitivity by focusing the sequencing efforts in specific regions of the genome (Mumbach et al., 2016, 2017). This is achieved using chromatin immunoprecipitation to capture genomic regions associated with a specific histone modification or protein of interest following Hi-C library preparation.

In our previous work, we used capture Hi-C to identify chromatin interactions at regions of the genome that contain variants associated with Ps in keratinocytes and skin derived CD8+ T cells (Ray-Jones et al., 2020). This targeted study suggested novel candidate causal genes for known Ps risk loci, such as the KLF4 gene at the intergenic locus 9q31. Here we use HiChIP to map active chromatin interactions genome wide on spontaneously immortalised keratinocytes (HaCaT) and a CD8+ T cell line derived from a cancerous skin plaque (MyLa). We show that this technique, in contrast with region capture Hi-C, shows significantly better enrichment for active regions of the genome. Moreover, it analyses interactions genome-wide, allowing us to discover candidate genes for a larger set of disorders and including more recently identified loci in our analysis.

We complement our dataset with matched RNA-seq and Hi-C to increase the information gained from these cell types. Because the disorders studied have a significant immune component we include in our analysis publicly available HiChIP data for Naïve CD4+ T cells and a B cell-like lymphoblastoid cell line (Mumbach et al., 2017) to describe the GWAS associated loci which might be mediated by these cell populations.

After identifying genes that are linked by chromatin interaction to GWAS loci we show that that these genes enrich for pathways that are highly relevant to the underlying disease mechanisms. Using the results generated by our analysis we provide a functional mechanism that drives disease susceptibility in some loci. More importantly, we present how our results can also inform the association of different genes to specific loci and provide example of these for 4 distinct Ps loci. This has the impact of updating our view on the underlying disease mechanisms of these traits, facilitating future studies and drug discovery.

## Results

### A compendium of activity and chromatin interactions in keratinocytes

Chromatin interactions that are specific for active regulatory elements such as enhancers and promoters, were identified using HiChIP through chromatin immunoprecipitation for the active histone mark H3K27ac. We analysed H3K27ac HiChIP data using a recently developed software specific for HiChIP data, FitHiChIP (Bhattacharyya, Chandra, Vijayanand, & Ay, 2019). This software identifies significantly enriched interactions with a much higher accuracy than previously available software. Interactions and peaks are highly reproducible and cell type specific, as shown by unsupervised clustering and principal component analysis (figure 1A-C). Because the samples cluster strongly by condition, we decided to merge the two replicates together to increase power as we noticed that the number of significant interactions reported by FitHiChIP depends on the read depth (Table S1). For example, in naïve HaCaTs the two replicates yielded 13950 and 38118 interactions, while the merged dataset yielded 58268 interactions.

**Figure 1.**
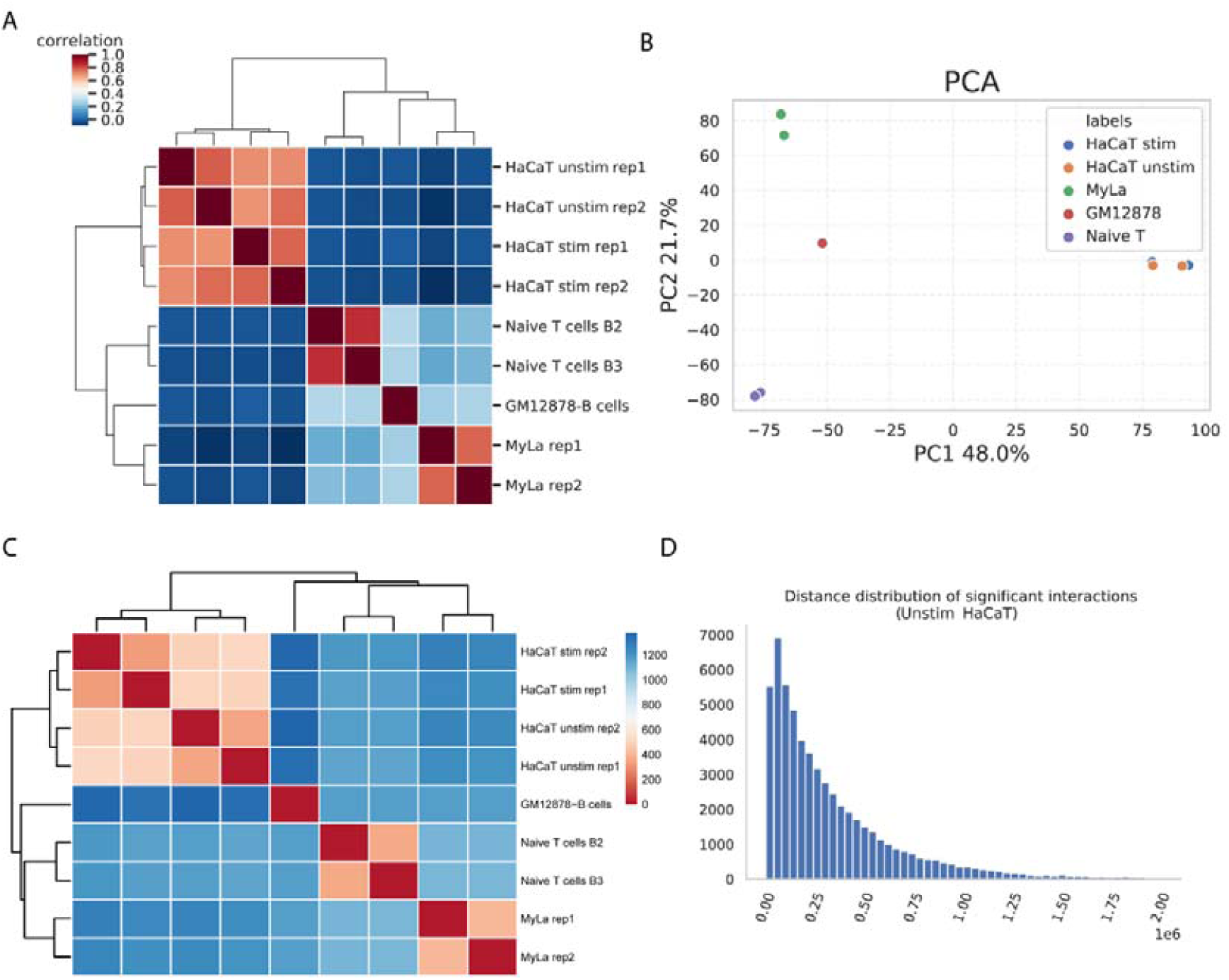
A) hierarchical clustering of the loops for the individual HiChIP samples using correlation. B) PCA of the loops for the individual HiChIP samples. C) hierarchical clustering of the HiChIP peaks for the individual samples using Euclidean distance. D) Distance distribution of the significant fithichip interactions for Naïve HaCaT HiChIP, a representative example.

We identified more than 50 000 significant interactions genome wide from each of our HiChIP datasets (summary statistics in table S2). The median interaction distance was 250 kb (figure 1D and S1A). These values are consistent with the results derived from the public datasets. Moreover, the vast majority (90%) of interactions were within topologically associating domains (TADs) identified from our Hi-C datasets derived from the same cell lines (figure 2C, summary statistics in table S2). This is consistent with recent reports about TADs determining the scope of gene regulation (Rao et al., 2014; Rowley & Corces, 2018).

**Figure 2.**
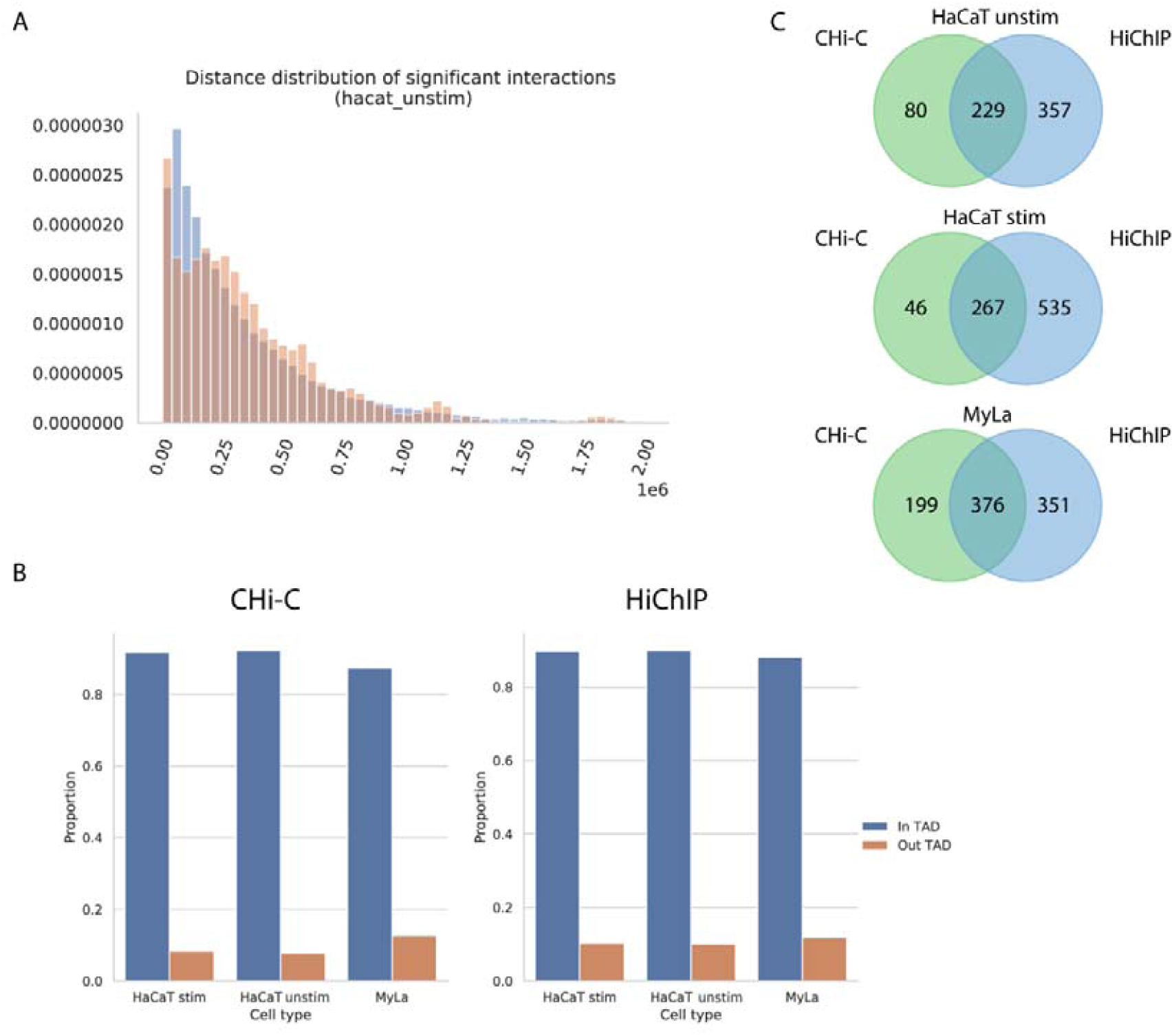
A) Distance distribution of the significant CHiCAGO interactions for unstimulated HaCaT region capture Hi-C (orange), overlapped over significant HiChIP interactions (blue). B) Proportion of significant interactions that are within TADs and that cross TAD boundaries for region capture Hi-C data and HiChIP data. C) Overlap of the genes identified through region capture Hi-C and HiChIP in HaCaT and MyLa cells for the same regions across all diseases studied.

Data from RNA-seq confirmed successful IFN-γ stimulation of the HaCaT cells, with 535 genes differentially expressed and clearly enriching for pathways related to IFN-γ stimulation (Supplementary figure S2). This has a significant impact on the number of interactions that originate from these genes: genes overexpressed upon stimulation have 7.08 interactions per gene in stimulated cells compared to 4.48 in unstimulated.

### HiChIP strongly enriches for interactions involving active regions of the genome in contrast with capture Hi-C

We compared this HiChIP data with our previous HaCaT and MyLa capture Hi-C data (Ray-Jones et al., 2020), which captured about 4,500 HindIII fragments containing GWAS SNPs for Ps, juvenile idiopathic arthritis, asthma, PsA, rheumatoid arthritis and SSc, curated from the largest available meta-analyses at the time and additional smaller GWAS.

Our previous capture Hi-C study identified about 35,000 significant interactions across all diseases studied, with a median interaction distance of 280 kb (figure 2A and S1B). Similar to HiChIP data, the majority of the interactions reside within TADs (Figure 2B). There is significant overlap between capture Hi-C and HiChIP (Figure 2C). For example, we recover the vast majority of the interacting genes highlighted in the previous work, such as *IL23A, STAT3, B3GNT2, COMMD1, ERRFI1* and *SOX4*.

Despite this, comparing HiChIP with Capture Hi-C we see that, although significant Capture Hi-C interactions are enriched for H3K27ac (figure S3), they are not specifically selected for active genes on enhancers. The majority of significant interactions (80% for unstimulated HaCaT) do not overlap H3K27ac peaks. This is in stark contrast with results from HiChIP for which, as expected, 99.8% of interactions overlap a H3K27ac peak at one end while 92.5% of interactions overlap a peak at both ends.

Moreover, Capture Hi-C significant interactions do not seem to be specific for active genes: 36% of the genes interacting with baits in unstimulated HaCaT are not expressed. This is in contrast with HiChIP, for which the majority of interactions are coming from active H3K27ac regions and 82% interacting genes from the same capture Hi-C regions were found to be expressed. This supports that HiChIP results are greatly enriched for active and informative interactions compared to Capture Hi-C. In addition, HiChIP finds significantly more interacting genes than Capture Hi-C (figure 2C). We note that whilst the Capture Hi-C findings and HiChIP findings are complementary, they enrich for different features and therefore represent different viewpoints. In addition, HiChIP allows for interpretation across different immune/skin-mediated diseases whereas targeted Capture Hi-C is biased by the selection of baits. Therefore, we focus on HiChIP-implicated genes for the remainder of this study.

### Chromatin contacts identified by HiChIP are confirmed by functional evidence

We first wanted to confirm that HiChIP can successfully discover important features involved in gene regulation. To do this we interrogated our HaCaT stimulation dataset and used RNA-seq to discover differentially expressed genes. We identify 535 differentially genes which are strongly enriched for interferon related pathways (figure S1) and assume that these genes are regulated through enhancers binding transcription factors related to response to the stimulation. As expected, H3K27ac peaks linked through HiChIP significant interactions to these genes are enriched for TF motifs that are known to be involved in the response to IFN-γ stimulation/viral infection (Ramezani, Nahad, & Faghihloo, 2018; Schroder, Hertzog, Ravasi, & Hume, 2004) and contain the similar motifs as those identified by differentially bound peaks (figure 3A).

**Figure 3.**
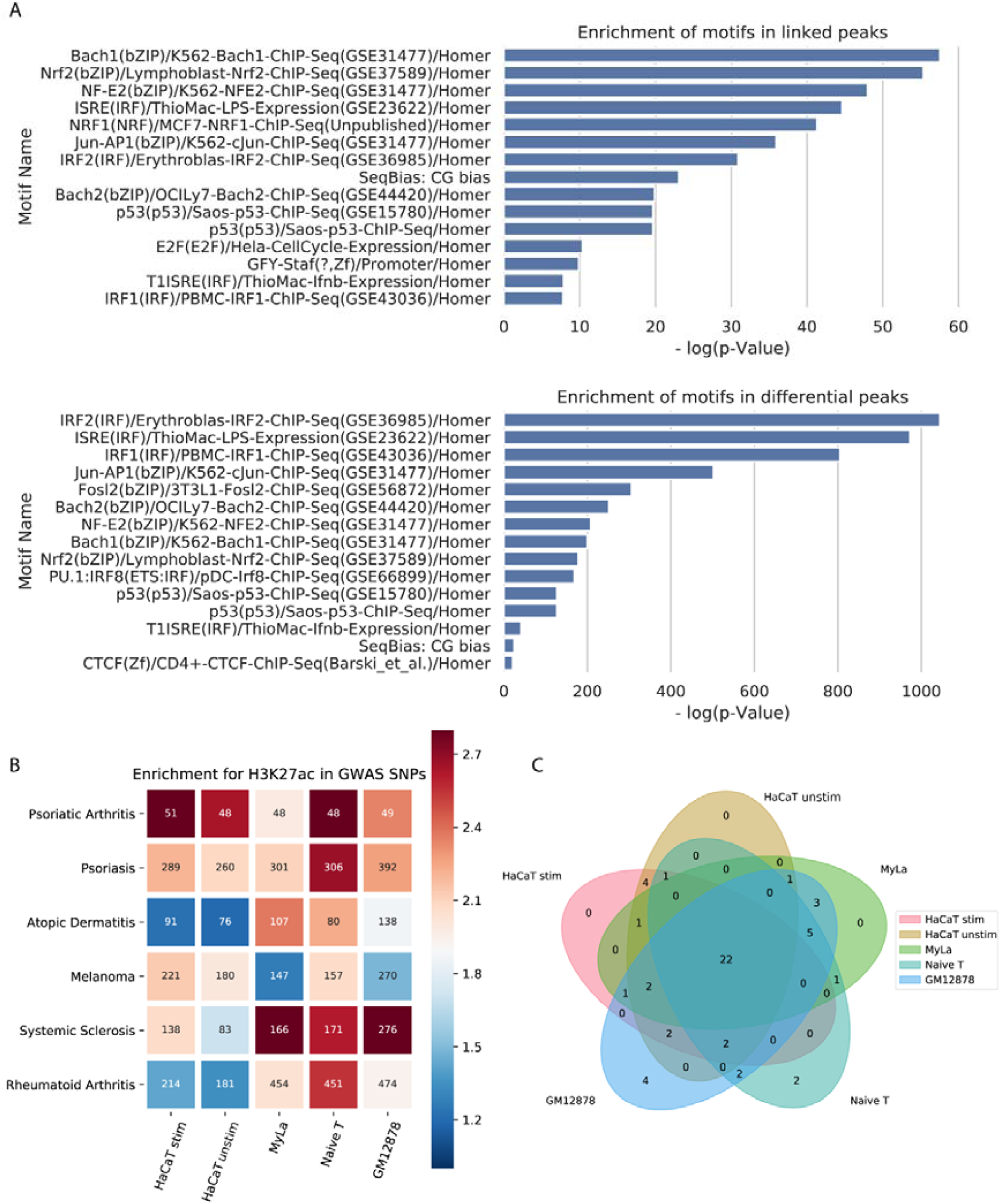
A) Pathways identified from differentially bound HiChIP H3K27ac peaks and HiChIP H3K27ac peaks linked to genes by HiChIP interactions to promoters of genes that were found to be differentially expressed during IFNγ stimulation. B) Heatmap showing the enrichment over background that disease associated variants have in HiChIP H3K27ac signal. Numbers inside each square represent the number of disease associated variants that directly overlap HiChIP H3K27ac peaks in each of the studied cell types. C) Venn diagram showing the number of psoriasis loci which have at least one variant overlapping a HiChIP H3K27ac peak in each cell type.

Next, we wanted to assess how HiChIP datasets can recapitulate results from large scale eQTL studies in the discovery of genes dysregulated by variants associated with disease. We used Ps loci as a proof of concept and identified the genes for which expression levels were influenced by GWAS SNPs. We tested our results from naïve T cells and GM12878 cells with the largest blood eQTL database available, eQTLgen (Võsa et al., 2018). Our HiChIP analysis showed 51% recall rate and 32% specificity compared to the genes identified from eQTLs. This shows very high concordance despite the fact that lymphocytes constitute a minority of cells present in whole blood.

We then compared our HaCaT keratinocyte datasets with the GTEX dataset from sun exposed skin (GTEx Consortium, 2013). Here the GTEX dataset contains relatively few statistically significant eQTLs compared to the eQTLgen database. Using our data, we see a recall rate of 53% and a specificity of 7.5%. Again, skin also contains more cell lineages than keratinocytes, and the low specificity is due to the lower power of this dataset compared to eQTLgen. Interestingly, 38 out of 52 skin specific eQTLs linked to Ps loci are recapitulated in the eQTLgen dataset.

These results demonstrate that our relatively small HiChIP dataset can successfully recapitulate an eQTL dataset produced with 31,000 samples and that we can discover significantly more genes compared to the GTEX database with more than 800 samples, giving us power to identify interactions in disease relevant cell types.

### GWAS variants are significantly enriched for active chromatin in disease relevant cell types

We wanted to test the assumption that GWAS results from various diseases would enrich for regulatory elements that are active in cell types relevant to their disease mechanism. To do so we used the H3K27ac signal collected from our HiChIP data using our recently developed tool, HiChIP-Peaks (Shi, Rattray, & Orozco, 2019), to estimate an enrichment score of these SNPs over a random background (figure 3B). As expected, SNPs associated with Ps, PsA, atopic dermatitis, melanoma and SSc show enrichment for the activity marker H3K27ac in HaCaT and MyLa cells. As a control we used RA, which is characterized by joint inflammation like PsA, but there is an absence of skin involvement. As expected, RA has much stronger enrichment in T cells and B cells relative to keratinocytes compared to the studied dermatological traits, such as Ps, PsA and melanoma. Interestingly PsA seems to have a much higher enrichment in keratinocytes than Ps even though most of the PsA loci are also shared by Ps. We think this reflects the fact that the PsA loci probably represent the strongest Ps signals as the PsA GWAS had a limited sample size.

The importance of studying disease specific cell types is evidenced by the fact that many SNPs overlap active regions of the genome only in specific cell types. Taking Ps as an example, we see that of the 45 loci that overlap a H3K27ac peak in any cell line, only 22 do so in all cell types, while 4 loci overlap a peak only in keratinocytes (figure 3C, other diseases in figure S4). Finally, for all traits studied, IFN-γ stimulated cells provided an increased enrichment of H3K27ac levels on the GWAS loci compared to their respective background. In Ps, 47 (15%) more SNPs are overlapping H3K27ac peaks in stimulated HaCaT compared to unstimulated providing potential involvement in a significant larger number of SNPs.

### Linking genes to disease associated loci provide functional dissection of disease mechanisms

Using the combined HiChIP datasets and cell-type matched RNA-seq data we identified transcribed genes whose transcription start sites (TSSs) either overlap or are linked by chromatin interaction to GWAS SNPs associated with PsA, Ps, atopic dermatitis, melanoma and SSc. We identify between 77 and 481 potentially linked genes for each disease studied (table 1, supplementary table 1). Importantly, these genes were strongly enriched for disease relevant pathways. For example, genes linked to melanoma loci were enriched for replicative senescence and cell cycle pathways, while genes linked to Ps and PsA were linked to cytokine signalling and IL-23 (figure 4), which is a pathway targeted by multiple novel treatments (Sakkas, Zafiriou, & Bogdanos, 2019).

**Table 1.** Number of genes identified associated for each disease studied.

| Disease | Number of genes identified |
| --- | --- |
| PsA | 77 |
| Ps | 374 |
| Eczema | 150 |
| Melanoma | 157 |
| Scleroderma | 171 |

**Figure 4.**
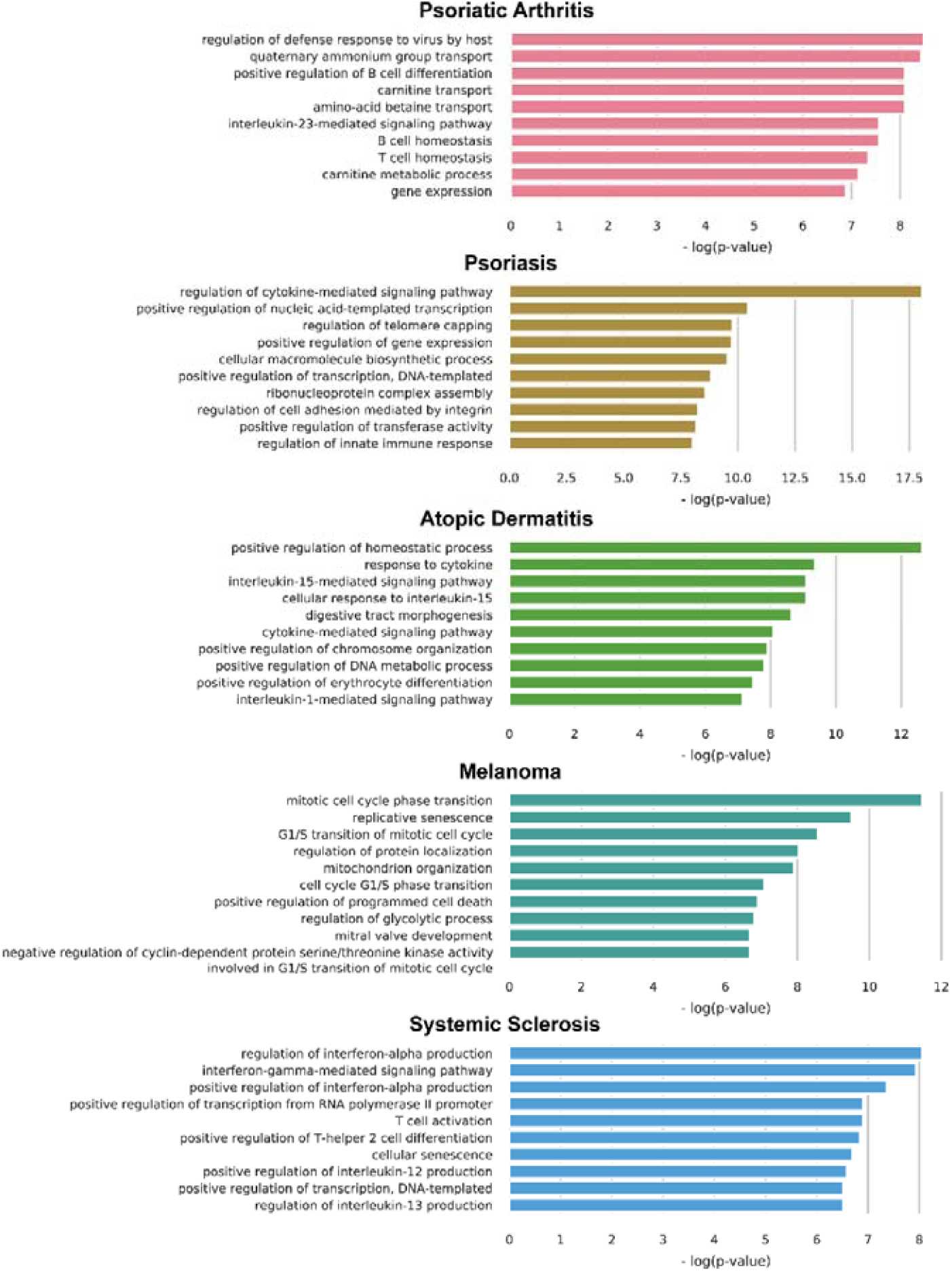
Top pathways enriched by the genes linked to disease associated loci from all cell types.

We found some differences in disease pathways that reflect the underlying pathology, for example, Ps pathways included regulation of cell adhesion mediated by integrin (important to the epithelium), that was not found in PsA: a related disorder that shares much disease background with Ps but primarily affects joints and not always skin.

Although some genes were common in all cell lines (e.g. in Ps 18.7%) most genes were specific to one or a few cell types (figure S5). Moreover, we found that most loci also implicate more than one gene, with on average 6.8 genes per loci in Ps. Interestingly the number of genes implicated by our method correlated significantly with the number of genes that were linked by eQTL for the same loci (p = 6.05e-14) (figure S6).

Using this data, we can provide functional insight into the mechanism involved in mediating disease susceptibility. For example, Ps and atopic dermatitis have distinct associations at the *ETS1* locus. However, whilst the Ps locus directly overlaps the promoter of *ETS1*, the association with atopic dermatitis is located 130 kb downstream of the gene (200 kb from the promoter). We observe significant chromatin interactions between the atopic dermatitis locus and the *ETS1* promoter in Naïve T cells, Myla cells and GM12878 cells (figure 5, S7A), while this enhancer was not active in HaCaT cells. Our data suggests a putative mechanism for how the distinct disease associations at this locus are mediated by a single gene.

**Figure 5.**
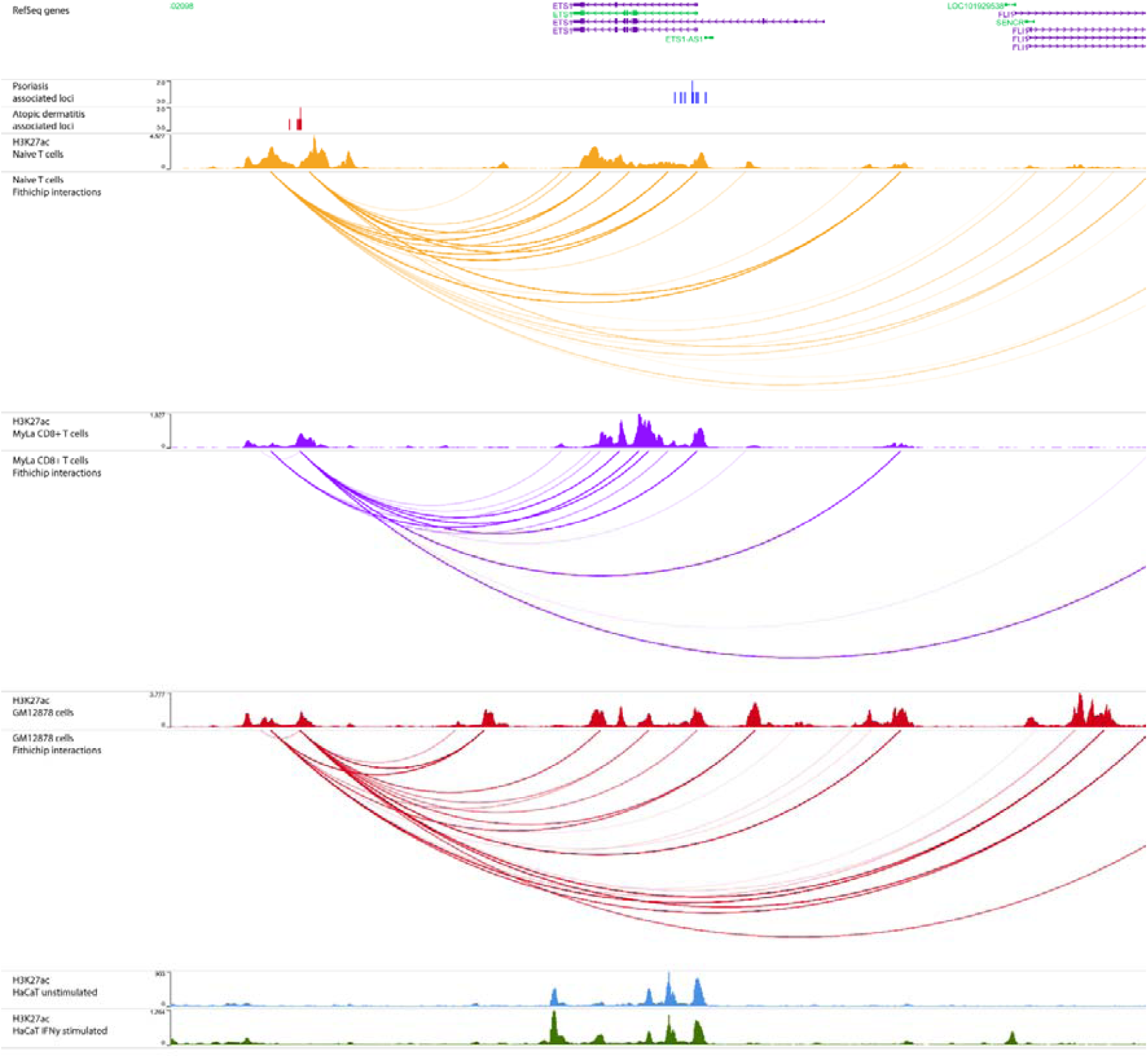
HiChIP interactions from the ETS1 locus provide a functional mechanism for this locus in atopic dermatitis. Tracks (in order): RefSeq genes; SNPs associated with Psoriasis (r^2^ > 0.8); SNPs associated with Atopic Dermititis (r^2^ > 0.8); H3K27ac signal in naïve T cells; Significant long range interactions originating from the atopic dermatitis associated locus in naïve T cells; H3K27ac signal in MyLa cells; Significant long range interactions originating from the atopic dermatitis associated locus in MyLa cells; H3K27ac signal in GM12878 cells; Significant long range interactions originating from the atopic dermatitis associated locus in GM12878 cells; H3K27ac signal in unstimulated HaCaT cells; H3K27ac signal in IFN-γ stimulated HaCaT cells.

### HiChIP identifies novel genes associated with psoriasis loci in a cell-type specific manner

The results from our analysis can be applied to augment existing understanding of disease associated loci and link novel genes or change the ones currently linked. For the purpose of this work we focus on 4 Ps associated loci which show cell type specific and previously unknown interactions.

The Ps locus indexed by SNP rs73178598 is located in an intergenic region overlapping an antisense RNA, *SATB1-AS1*. We found a 240 kb T cell specific interaction present in Naïve T cells and MyLa cells but not keratinocytes linking this locus with the promoter of SATB1 (figure 6). Interestingly, upon examination this is also visible in Hi-C maps (figure 6s7b). This gene has not been previously linked to Ps genetically. Silencing of SATB1 has been shown to have a similar effect to IFN-γ stimulation on MHC chromatin organization (Pavan Kumar et al., 2007) and is known to be an important regulator of regulatory T cells and autoimmunity (Beyer et al., 2011).

**Figure 6.**
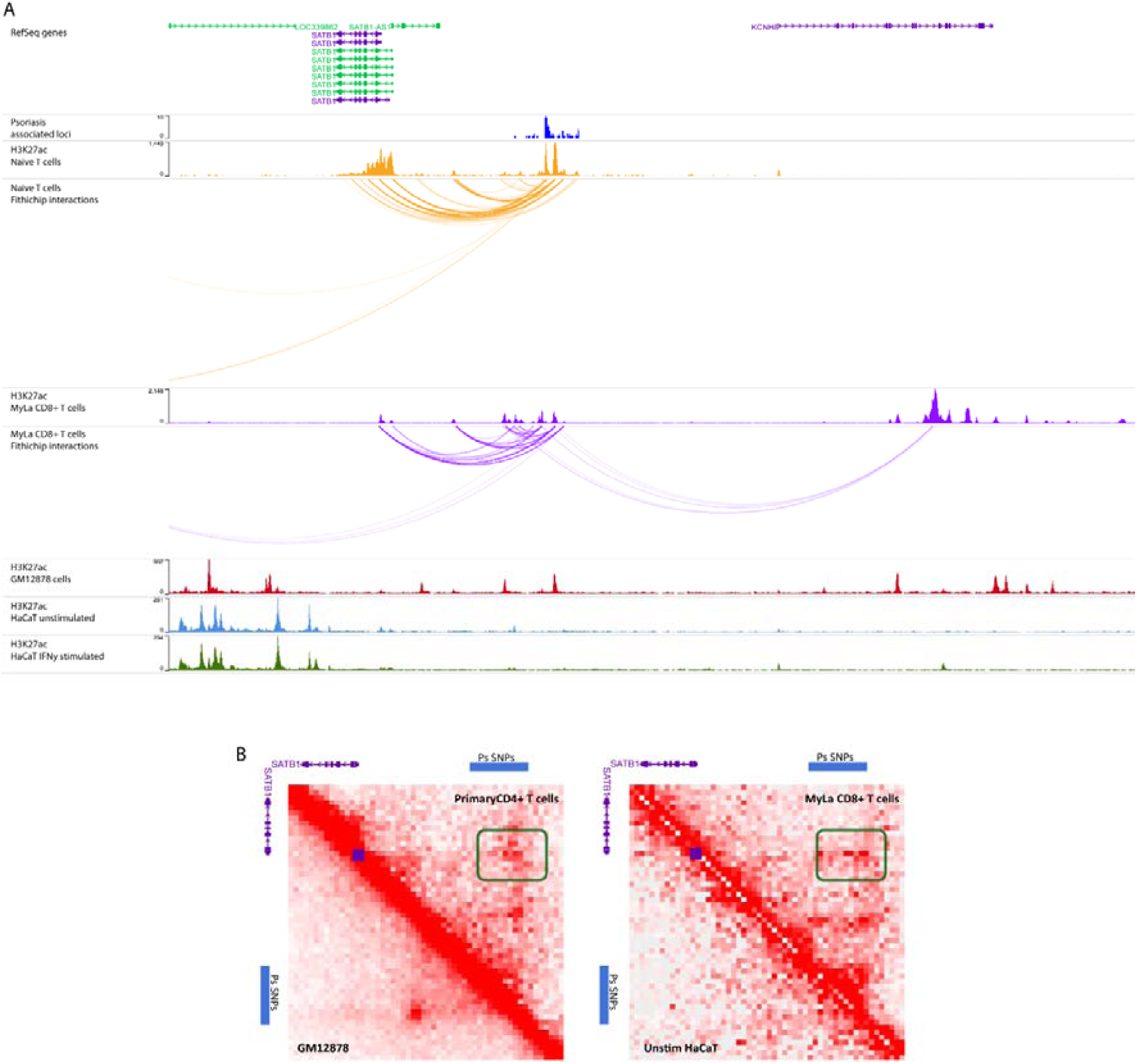
A): HiChIP interactions from the *SATB1/KCNH8* locus link *SATB1* as the target gene in this locus. Tracks (in order): RefSeq genes; SNPs associated with Psoriasis (r^2^ > 0.8); H3K27ac signal in naïve T cells; Significant long range interactions originating from the psoriasis associated locus in naïve T cells; H3K27ac signal in MyLa cells; Significant long range interactions originating from the psoriasis associated locus in MyLa cells; H3K27ac signal in GM12878 cells; H3K27ac signal in IFN-γ stimulated HaCaT cells; H3K27ac signal in naïve T cells. B): Hi-C contact maps show a cell type specific loop (highlighted in green) that is present between the psoriasis SNPs and the *SATB1* promoter (purple square).

The Ps loci indexed by SNP rs9504361 is intronic to *EXOC2* and is typically associated with this gene. However, despite *EXOC2* being widely expressed and its expression being associated with this SNP in blood, H3K27ac occupancy is specific to MyLa in our analysis. A long-range interaction was detected between this SNP and the promoters of *IRF4* and *DUSP22*, specifically in this cell type and not in Naïve CD4^+^ T cells or HaCaT keratinocytes (figure 7, S7B). *IRF4* is a lymphocyte specific transcription factor that negatively regulates Toll-like-receptor (TLR) signalling, as part of the interferon response, central to the activation of innate and adaptive immune systems (Huber & Lohoff, 2014). It was found to be overexpressed in psoriatic skin lesions (A. Ni et al., 2012). DUSP22 is a phosphatase that might be involved in the JNK signalling pathway and has been shown to be associated with lymphomas (Paydas, Bagir, Ergin, Seydaoglu, & Boz, 2019; Zeke, Misheva, Reményi, & Bogoyevitch, 2016).

**Figure 7.**
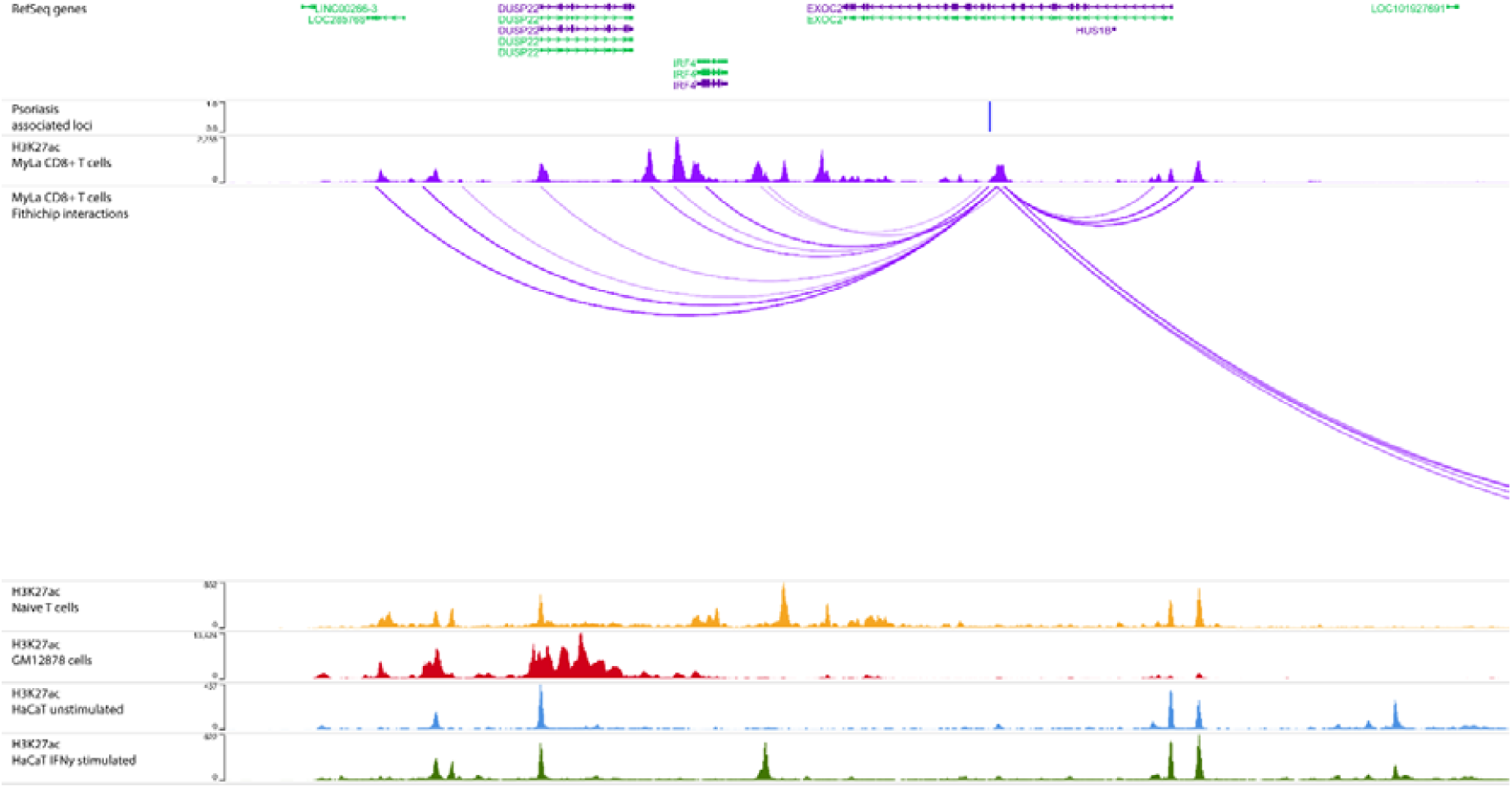
HiChIP interactions from the *EXOC2/IRF4/DUSP22* locus link *IRF4* and *DUSP22* as candidate genes in psoriasis. Tracks (in order): RefSeq genes; SNPs associated with Psoriasis (r^2^ > 0.8); H3K27ac signal in naïve T cells; H3K27ac signal in MyLa cells; Significant long range interactions originating from the atopic dermatitis associated locus in MyLa cells; H3K27ac signal in naïve T cells; H3K27ac signal in GM12878 cells; H3K27ac signal in unstimulated HaCaT cells; H3K27ac signal in IFN-γ stimulated HaCaT cells.

Another example of a novel gene target can be found at the 1p36 locus (rs10794648), which to-date has been associated with *IFNLR1* (also known as IL28-RA) as it is the closest gene to the associated SNPs (Strange et al., 2010; Stuart et al., 2015). *IFNLR1* protein encodes part of a receptor for IFN-γ that is presented in the epidermis, and thought to promote antiviral response in Ps (Lazear, Nice, & Diamond, 2015). However, our HiChIP data showed long-range interactions between the Ps SNPs at 1p36 and the distal gene *GRHL3* primarily in HaCaT keratinocytes (figures 8, S8A, S9). This gene encodes a transcription factor, upregulated in psoriatic lesions and required for repair of the epidermal skin barrier following immune-mediated injury (Gordon et al., 2014). Targets of the GRHL3 transcription factor include further GWAS-implicated genes *IVL* (involved in keratinocyte differentiation) (Watt, 1983) and *KLF4* (transcription factor involved in keratinocyte differentiation and skin barrier formation). *KLF4* was shown previously as upregulated by IFN-γ and implicated as a likely functional target gene in the 9q31 locus through chromatin looping and CRISPR activation (Ray-Jones et al., 2020).

**Figure 8.**
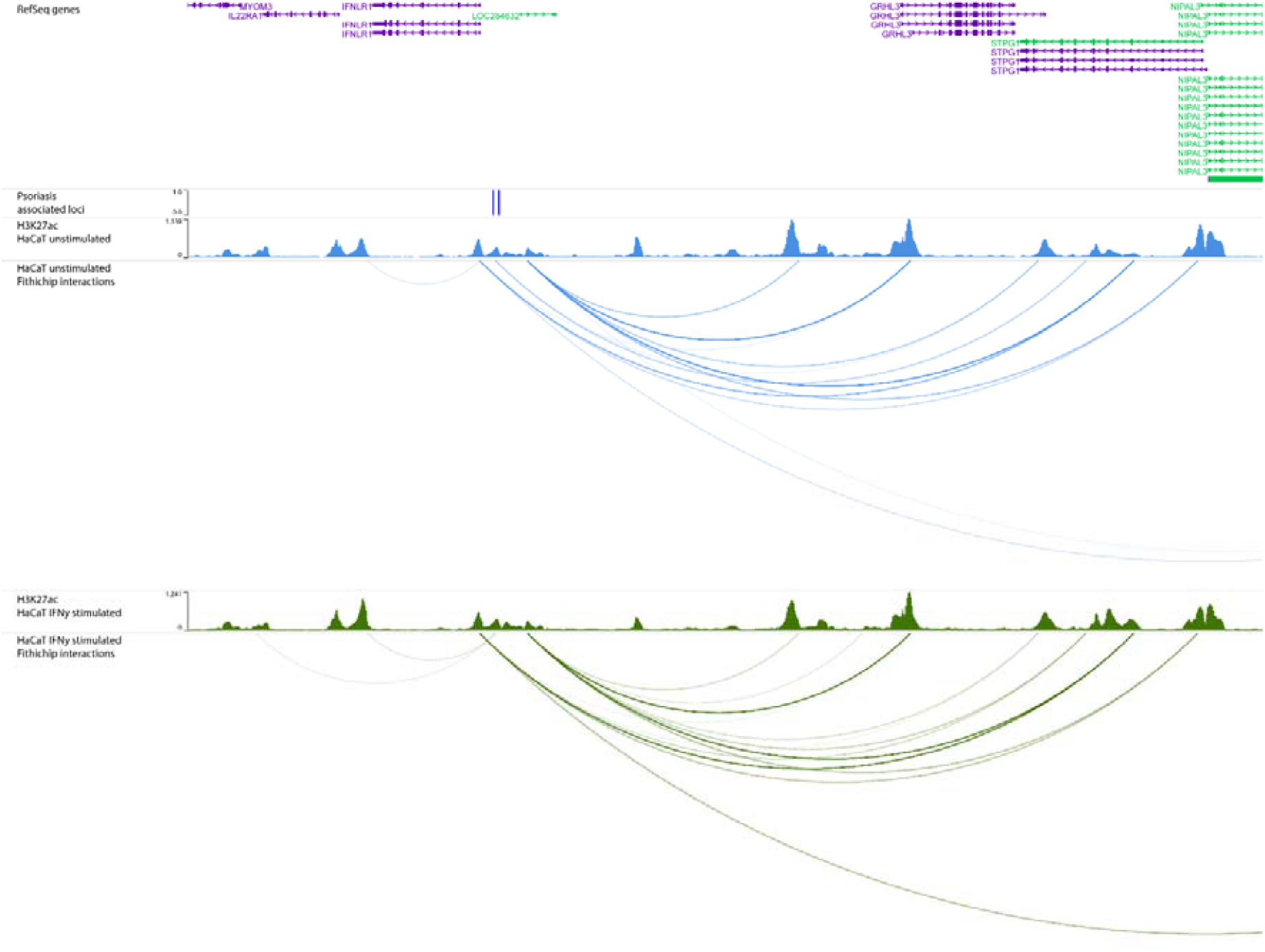
HiChIP interactions from the *IFNLR1/GRHL3* locus link *GRHL3* as a candidate gene in psoriasis. Tracks (in order): RefSeq genes; SNPs associated with Psoriasis (r^2^ > 0.8); H3K27ac signal in unstimulated HaCaT cells; Significant long range interactions originating from the psoriasis associated locus in unstimulated HaCaT cells; H3K27ac signal in IFN-γ stimulated HaCaT cells; Significant long range interactions originating from the psoriasis associated locus in IFN-γ stimulated HaCaT cells.

Finally, the locus indexed by the Ps SNP rs73183592 with previously unknown function was linked via a long-range interaction spanning about 500 kb to *FOXO1* (figure 9, S8B), a gene with important functions in regulatory T cells. Interestingly, this interaction was identified only in HaCaT keratinocytes in our analysis. Dysregulation of this pathway was also found to be important in the development of Ps (Li et al., 2019; Zhang & Zhang, 2019).

**Figure 9.**
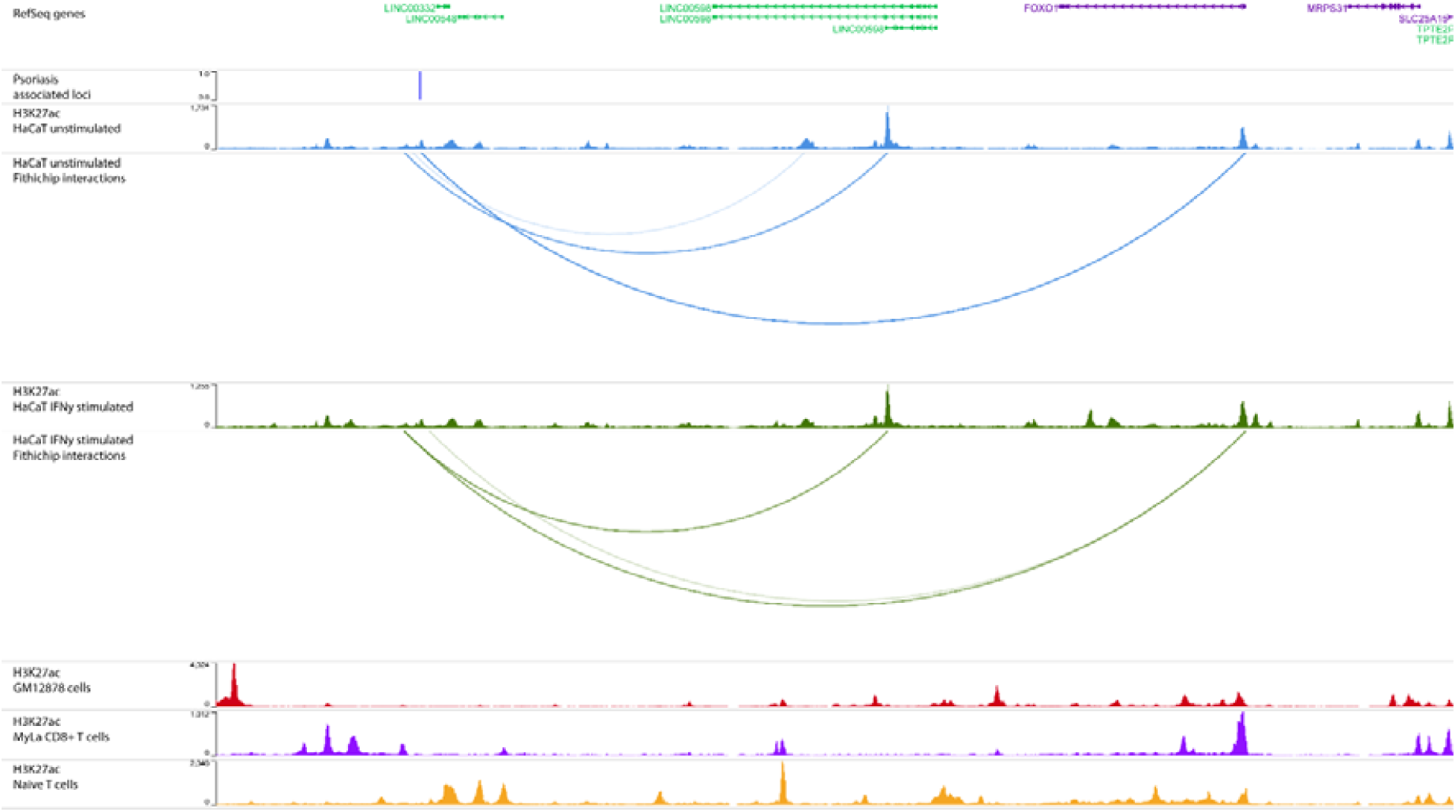
HiChIP interactions from the *FOXO1* locus link *FOXO1* as a candidate gene in psoriasis. Tracks (in order): RefSeq genes; SNPs associated with Psoriasis (r^2^ > 0.8); H3K27ac signal in unstimulated HaCaT cells; Significant long range interactions originating from the psoriasis associated locus in unstimulated HaCaT cells; H3K27ac signal in IFN-γ stimulated HaCaT cells; Significant long range interactions originating from the psoriasis associated locus in IFN-γ stimulated HaCaT cells; H3K27ac signal in GM12878 cells; H3K27ac signal in MyLa cells; H3K27ac signal in naïve T cells.

## Discussion

Chromatin conformation and functional genomics studies have the potential to uncover the underlying mechanisms that drive the disease susceptibility of many complex traits. Although these techniques are very promising, there has been a lack of studies in disease relevant cell types, and as recently evidenced, both chromatin interactions and gene regulation are cell type and stimulation specific (Burren et al., 2017; Dixon et al., 2012; Hansen et al., 2018; Mumbach et al., 2017; Rao et al., 2014; Rubin et al., 2017; Schmitt et al., 2016; Siersbæk et al., 2017).

Here we used H3K27ac HiChIP, a novel technique that allows combined analysis of both chromatin conformation and chromatin activity, to create the first global study of promoter-enhancer interactions in keratinocytes and tissue resident CD8^+^ T cell lines. Equivalent data has so far only been generated in immune cells with few examples in other cell populations, such as HiChIP in endometrial cancer cells (O’Mara, Spurdle, & Glubb, 2019), and promoter-capture Hi-C in neuronal cells (Song et al., 2019), cardiomyocytes (Choy et al., 2018) and pancreatic islets (Miguel-Escalada et al., 2019). Interestingly, an increased enrichment of H3K27ac peaks over GWAS loci in our data suggests that keratinocytes stimulated with IFN-γ stimulation may be more relevant than naïve keratinocytes for all diseases studied. A common limitation of these studies (including this one) is that they make use of immortalized cell lines or differentiated pluripotent stem cells. Ideally, future studies would involve primary cells or tissue from patients.

We have analysed the effectiveness of these techniques to link functional relevant elements and shown how we can use them to study chromatin interaction and activity in a cell type specific manner. With the combination of public data for cell types relevant in the disorders studied we have analysed the potential information that can be gained for a number of skin-related disorders.

Using these datasets, we study all disease associated SNPs for PsA, Ps, atopic dermatitis, melanoma and SSc, and identify all the genes that are linked by chromatin interactions to these variants. We show that these genes enrich for disease relevant pathways and provide tables for all loci (supplementary tables). We demonstrate how this data allows us to identify novel mechanisms and we show, using four distinct Ps associated loci, how we can provide functional insight and link novel genes or change the ones currently linked.

We highlight two loci in which keratinocytes provide us cell type specific interactions that were not found in the more commonly used immune cell populations. We link the locus indexed by rs73183592 to *FOXO1*, a gene located about 500 kb away from the locus. We also show a link between the Ps SNP rs10794648 and *GRHL3*, a gene involved in the epidermis and fundamental to keratinocytes’ function. Phenotypically, Ps, is a disease primarily affecting skin barrier and this locus possibly explain some of the genetic background that leads to developing the disease. These novel connections have the potential to be used as therapeutic targets in drug repurposing and discovery, as recently applied for other diseases (Fang et al., 2019; P. Martin et al., 2019). As a proof of concept, we tested the genes identified in this work by querying the drugbank database (Wishart et al., 2008) in search for drugs that are currently used and ones that could be repurposed. Across the diseases studied we identified 192 genes that are targeted by approved drugs in some diseases, corresponding to 299 drugs that could be potentially repurposed (supplementary table 2).

Whilst these results are important, a limitation of our study is that further biological and functional validation is required, such as those included in our recent study of the *KLF4* locus (Ray-Jones et al., 2020). Nevertheless, through our analysis we present a list of potential target genes and pathways for mediating disease risk for five complex diseases and by documenting individual loci we highlight the mechanisms by which this risk is mediated. The genes and mechanisms represent a useful resource for further research aimed at characterising how increased disease susceptibility is mediated for these and other complex diseases.

## Methods

### Cell culture

HaCaT keratinocyte cells were obtained from Addexbio (T0020001); these are in vitro spontaneously transformed keratinocytes from histologically normal skin. Cells were cultured in high-glucose Dulbecco’s modified eagle’s medium (DMEM) supplemented with 10% foetal bovine serum (FBS) and penicillin-streptomycin (Thermo Fisher Scientific, final concentration 100 U penicillin, 0.1 mg streptomycin/ml). For HaCaT stimulation experiments, the media was supplemented with 100 ng/mL recombinant human IFN-γ (285-IF-100; R&D Systems) and cells incubated for 8 hours prior to harvest.

My-La CD8+ cells were obtained from Sigma-Aldrich (95051033). These cells are cancerous human T-lymphocytes derived from a patient with mycosis fungoides. Cells were cultured in Roswell Park Memorial Institute (RPMI) 1640 medium supplemented with 10% AB human serum (Sigma Aldrich), 100 U/mL recombinant human IL-2 (Sigma-Aldrich) and penicillin-streptomycin (final concentration 100 U penicillin, 0.1 mg streptomycin/ml).

### Cell crosslinking for chromatin-based experiments

HaCaT cells were crosslinked for 10 minutes in 1% formaldehyde and the reaction was quenched with 0.135M glycine. The crosslinked cells were scraped from the flask, pelleted, washed in PBS and the supernatant removed. My-La cells were crosslinked for 10 minutes in 1% (HiChIP) or 2% (Hi-C) formaldehyde and the reaction was quenched with 0.135M glycine and the supernatant removed. Fixed cells were snap frozen on dry ice and stored at −80°C.

### HiChIP library generation and processing

HiChIP libraries were generated according to the Chang Lab protocol (Mumbach et al., 2016). 10 million crosslinked cells were lysed and the chromatin digested using 375 U of MboI (NEB, R0147M) for 4 hours at 37°C. Fragment ends were filled in using dCTP, dGTP, dTTP and biotin-14 dATP (Life Technologies) and ligated at room temperature overnight. The nuclei were lysed and the chromatin sheared to lengths of approximately 200-700 bp using a Covaris S220. Immunoprecipitation was performed overnight at 4°C using 20 µg of H3K27ac antibody (Abcam ab4729). The DNA was captured on a 1:1 mixture of protein A and G Dynabeads (Invitrogen 10001D and 10003D). After washes, the DNA was eluted with proteinase K at 65°C overnight. The sample was cleaned using Zymo Clean and Concentrator Columns (Zymo D4013) and quantified using the Qubit DNA HS kit. 20-35 ng of DNA was taken forward for biotin-pulldown with streptavidin C-1 beads at room temperature for 30 minutes. The beads were suspended in TD buffer from the Nextera kit and transposed with Tn5 (Illumina) at 55°C for exactly 10 minutes. The volume of Tn5 was dependent on DNA quantity and defined by the original HiChIP protocol. After washes, the library was amplified off the beads using Phusion polymerase and Nextera indexing primers (Illumina). AMPure XP beads (Beckman Coulter, A63882) were used to select fragments approximately 300 – 700 bp in length. Quantification and quality control of the final HiChIP library was conducted using a Bioanalyzer and KAPA quantification kit (Kapa Biosystems). Libraries underwent Next Generation Sequencing on a HiSeq 2500 generating 100 bp paired-ends.

Sequencing data for the HiChIP libraries was filtered and the adapters were removed using fastp v0.19.4 (S. Chen, Zhou, Chen, & Gu, 2018). The reads were then mapped to the GRCh38 genome with Hi-C Pro v2.11.0 (Servant et al., 2015), using default settings. Enriched regions (H3K27ac peaks) were identified using HiChIP-peaks v 0.1.1 (Shi et al., 2019) with default settings and FDR < 0.01. Loops were identified using FitHiChIP (Bhattacharyya et al., 2019) using the following settings: Coverage normalization, stringent background with merging enabled, peaks generated from HiChIP-peaks and 5 kb bin size. Viewpoints for the figures were generated by selecting interactions originating from within 10 kb of the index SNP.

### HiChIP clustering and principal component analysis

To validate the reproducibility and cell type specificity of our HiChIP loops we collected the top 10000 significant loops from each individual replicate and combined it to create a set of 82545 loops across all samples. For each of these loops we then collected the raw FitHiChIP p-value from each sample from the raw interactions file (replacing missing entries with 1). We then ran hierarchical clustering using correlation (seaborn clustermap) and PCA (Scikit-learn) analysis on the resulting data matrix.

For the clustering of the peaks we used the included differential peak calling module of HiChIP-peaks (Shi et al., 2019) and used the data matrix provided to run hierarchical clustering using Euclidean distance.

### Hi-C library generation and processing

In-situ Hi-C libraries were generated for HaCaT and MyLa cell lines as previously described (Ray-Jones et al., 2020). 50 million crosslinked cells were lysed and the chromatin digested with HindIII at 37°C overnight. Restriction cut sites were filled in using dCTP, dGTP, dTTP and biotin-14-dATP (Life Technologies), then in-nucleus ligation was carried out at 16°C for 4-6 hours. Crosslinks were reversed by proteinase-K overnight at 65°C and RNA was digested using RNaseA for 60 minutes at 37°C. The DNA was purified by sequential phenol and phenol-chloroform extractions and ethanol-precipitated at −20°C overnight, followed by two further phenol-chloroform extractions and a second overnight precipitation.

A 40 µg aliquot of DNA was taken forward for further processing following QC steps. T4 DNA polymerase used to remove biotin-14-dATP from non-ligated ends then the DNA purified by phenol-chloroform extraction and ethanol precipitation overnight. The DNA was sheared using a Covaris S220 sonicator and end-repair was performed using T4 DNA polymerase, T4 DNA polynucleotide kinase and DNA polymerase I, large (Klenow) fragment. The sample was purified using Qiagen MinElute Kit, with a modified protocol described by (Belton et al., 2012). Klenow (exo-) was used to adenylate DNA fragment ends and a double-sided SPRI bead size selection was used to obtain fragments of approximately 200-600 bp. Dynabeads MyOne Streptavidin C1 beads (Life Technologies) were used to pull down biotinylated fragments, which were then ligated to annealed Illumina sequencing adapters. PCR was performed using Phusion HF (NEB) and TruPE PCR primers (Illumina), then the amplified DNA was cleaned twice using 1.8X volume of SPRI beads. The quality and quantity of the Hi-C libraries was tested by Bioanalyzer and KAPA qPCR. Hi-C libraries were analysed by Next Generation Sequencing. The My-La Hi-C library was sequenced on an Illumina HiSeq 2500 generating 100bp paired ends. The HaCaT Hi-C libraries were sequenced on an Illumina HiSeq 4000 generating 75 bp paired ends.

The sequencing data was filtered and adapters were removed using fastp v0.19.4 (S. Chen et al., 2018). The reads were then mapped to the GRCh38 genome with Hi-C Pro v2.11.0 (Servant et al., 2015), using default settings. The Hi-C interaction matrices were normalised within Hi-C Pro using iterative correction and eigenvector decomposition (ICE). TADs were identified using OnTAD v1.2 (An et al., 2019), a novel Optimized Nested TAD caller for Hi-C data, using Hi-C data binned at a 40 kb resolution and a maximum TAD size of 4 mb. Files for visualisation were created using the hicpro2juicebox.sh utility and visualised in Juicebox (Durand et al., 2016). Maps were normalized with the balancing algorithm whenever that converged or the coverage (sqrt) method otherwise.

For primary Naïve T cells, we used a different protocol to generate the Hi-C maps. PBMCs were isolated from a buffy coat obtained from the National Blood Transfusion Service using a ficoll gradient. T-cells were isolated using an EasySep T-cell isolation kit according to the manufacturer’s instructions. 3 million cells were fixed with 2% formaldehyde for 10 minutes and then snap frozen. Hi-C libraries were generated using the Arima Hi-C kit following manufacturer’s instructions. Data was then processed and analysed in the same way as the other libraries.

For GM12878 cells we obtained 1.2B raw reads from Arima Genomics. This library was chosen because it was more directly comparable with our primary Naïve T cells library and, as it was generated using the Arima Hi-C protocol. Data was then processed and analysed in the same way as the other libraries.

### Region Capture Hi-C and overlap with HiChIP results

Region capture Hi-C libraries were generated as previously described from the Hi-C libraries as part of a previous study (Ray-Jones et al., 2020).

First, amplified Hi-C libraries were generated as above. Then, Hi-C DNA up to 750 ng was concentrated using a vacuum concentrator and bound to the capture baits in a single hybridisation reaction using SureSelectXT reagents and protocol (Agilent Technologies). The biotinylated baits were captured using Dynabeads MyOne Streptavidin T1 beads (Life Technologies). Following washes, the libraries were amplified on the beads using Phusion HF and barcoded TruPE primers then the amplified DNA cleaned twice using 1.8X volume of SPRI beads. The quality and quantity of the capture Hi-C libraries was tested by Bioanalyzer and KAPA qPCR (Kapa Biosystems). Capture Hi-C libraries were analysed by 75 bp paired-end Next Generation Sequencing on an Illumina NextSeq500 (My-La) or HiSeq 4000 (HaCaT). Capture Hi-C sequence data was quality filtered with fastp v 0.19.4 (S. Chen et al., 2018) and then processed through the Hi-C User Pipeline (HiCUP) v0.7.2 (Wingett et al., 2015) and mapped to the GRCh38 genome. For each cell type, the two biological replicates were simultaneously run through Capture Hi-C Analysis of Genomic Organisation (CHiCAGO) v1.10.1 (Cairns et al., 2016) in R v3.5.1 and significant interactions were called with a score threshold of 5. Enrichment for H3K27ac was calculated using the integrated tool in the CHiCAGO package with default settings and the H3K27ac peaks generated from the HiChIP data using HiChIP-Peaks.

To identify active enhancer-promoter interactions we kept the CHiCAGO interactions that originated from HiChIP H3K27ac peaks in the matching cell type. We then identified the expressed promoters (TPM>1) that were within 5 kb of the other end of the interactions.

To compare the results from Capture Hi-C with our new HiChIP libraries we determined the interactions that originated from within 5 kb of those 4500 capture fragments. Genes were then identified as described in the “Linking GWAS results to putative gene targets” section.

### RNA-seq

3’ mRNA sequencing libraries were generated for cell lines using the Lexogen QuantSeq 3’ mRNA-Seq Library Prep Kit FWD for Illumina. Libraries were sequenced using single-end Illumina SBS technology. Reads were quality trimmed using Trimmomatic v0.38 (Bolger, Lohse, & Usadel, 2014) using a sliding window of 5 with a mean minimum quality of 20. Adapters and poly A/poly G tails were removed using Cutadapt v1.18 (M. Martin, 2011) and then UMIs were extracted from the 5’ of the reads using UMI-tools v0.5.5 (Smith, Heger, & Sudbery, 2017). Reads were then mapped using STAR v2.5.3a (Dobin et al., 2013) on the GRCh38 genome with GENCODE annotation v29 (Harrow et al., 2012). Reads were de-duplicated using UMIs with UMI-tools and then counted using HTSeq v0.11.2 (Anders, Pyl, & Huber, 2015). Count matrixes were analysed in R 3.5.1 and normalisation and differential expression analysis was conducted using DESeq2 v1.22.2 (Love, Huber, & Anders, 2014). Differentially expressed genes were called with an adjusted P value of 0.10 (FDR 10%). For detection of expressed genes in the cell lines, we considered RNA-seq counts greater than 1 count per million.

### Public RNA-seq data

Public RNA-seq for the CD4 naïve t cell type was downloaded from (Bonnal et al., 2015). Accession ID: ERP004883. Raw sequencing reads were filtered and adapters and polyAs trimmed with fastp v 0.19.4 (S. Chen et al., 2018). Reads were then mapped with salmon v0.14.1 (Patro, Duggal, Love, Irizarry, & Kingsford, 2017) to the GRCh38 genome with GENCODE annotation v29 (Harrow et al., 2012).

TPM values were used later in the analysis for gene filtering.

### GWAS data

Genome wide significant (p-value < 5×10^-8) GWAS loci were downloaded for the following diseases: PsA (Bowes et al., 2015; Stuart et al., 2015), Ps (Tsoi et al., 2017), melanoma (Duffy et al., 2018), SSc (López-Isac et al., 2019), atopic dermatitis (Paternoster et al., 2015) and rheumatoid arthritis (Okada et al., 2014).

SNPs in high linkage disequilibrium (R^2 > 0.8) with the lead SNPs were identified using plink v1.90b3.39 on the 1000 genomes data v3 with population set to EUR.

### SNP enrichment

We obtained the H3K27ac signal tracks for each cell type from the HiChIP data using HiChIP-Peaks. This track corresponds to the signal for this marker of activity along the genome. We then calculated the median intensity of the signal over every SNP outside of the MHC and compared it with the median for a set of 1 million randomly generated positions to get an estimate of a genomic background for the signal to calculate an enrichment. We also calculated the number of individual SNPs that are located within a H3K27ac peak for each cell type.

### Linking stimulation responsive genes to enhancers

We identified the genes that were differentially expressed during the INF-γ stimulation as described previously. We then identified the promoter regions for these genes and identified the genomic regions that interacted with these promoters in any of the two conditions in HaCaT cells.

We intersected these regions with the H3K27ac peaks identified by HiChIP-Peaks to narrow down the exact location of the interacting enhancers and ran Motif enrichment analysis using HOMER v 4.8.3 (Heinz et al., 2010) with the findMotifsGenome.pl command and “–size given” parameter.

### Linking GWAS loci to putative gene targets

To identify the genes that were linked to disease associated SNPs we first identified the transcription start sites of all protein coding transcripts in the hg38 Gencode V29 annotation (Harrow et al., 2012). We associated all transcripts for which a TSS was located within 5 kb of a loop. We also associated the transcripts for which the TSS was within 1 kb of a SNP overlapping a H3K27ac peak as identified from HiChIP data.

All transcripts were then grouped by originating gene and all analysis was done at the gene level.

Lastly genes were filtered by expression level of at least 1 TPM in the corresponding cell line.

### Overlap with eQTL

We downloaded the full cis-eQTL datasets from the sun-exposed skin GTEx v7 dataset (GTEx Consortium, 2013) and the eQTLgen (version 2018/10/17) dataset (Võsa et al., 2018). To identify the eQTLs that were originating from the GWAS loci we simply queried every SNP that was in LD with the lead SNP and recorded all the genes that were significantly linked to those variants. Genes were filtered by expression TPM > 1 in our cell types. We then identified all the genes that are linked for a specific disease (in this example psoriasis) and compared this list with the list of genes that were identified from the same SNPs using the HiChIP interactions.

### Pathway analysis

The most enriched pathways for each disease were identified using the EnrichR (E. Y. Chen et al., 2013) web API with the gene set library set to GO_Biological_Process_2018. The pathways were then sorted by p-value and the top 10 enriched pathways were plotted.

### Drug discovery

Drug discovery was executed by querying the drugbank v5.1.4 database (Wishart et al., 2008). New drugs available for repurposing were identified by the approved tag in at least one disease and the target being one of the genes studied.

## Acknowledgements

The authors would like to acknowledge the assistance given by IT Services and the use of the Computational Shared Facility at The University of Manchester.

This work was funded by the Wellcome Trust (award references 207491/Z/17/Z and 215207/Z/19/Z), Versus Arthritis (award reference 21754), NIHR Manchester Biomedical Research Centre and the Medical Research Council (award reference MR/N00017X/1).

## Supplementary Tables and figures

**An active chromatin interactome elucidates the biological mechanisms underlying genetic risk factors of dermatological conditions in disease relevant cell lines**

**Table S1.** Number of interactions per replicate for the conditions.

|  | Replicate 1 | Replicate 2 | Combined |
| --- | --- | --- | --- |
| HaCaT unstim | 13950 | 38118 | 58268 |
| HaCaT stim | 14456 | 47100 | 68887 |
| MyLa | 33257 | 13743 | 51274 |
| Naïve T (Mumbach) | 29352 | 28405 | 42710 |

**Table S2.** Summary statistics of the datasets used in this project.

| Cell Type (HiChIP) | Valid interaction pairs (millions) | Median loop distance (kb) | Number of significant loops |
| --- | --- | --- | --- |
| HaCaT unstim (Keratinocytes) | 273.1 | 220 | 58268 |
| HaCaT IFN-γ stimulated | 309.8 | 235 | 68887 |
| MyLa (CD8+ T cells) | 288.6 | 300 | 51274 |
| CD4+ Naïve T cells | 167.8 | 310 | 42710 |
| GM12878 (B-cell) | 241.3 | 185 | 76451 |

| Cell Type (Hi-C) | Valid interaction pairs (millions) |
| --- | --- |
| HaCaT unstim (Keratinocytes) | 106.2 |
| HaCaT IFN-γ stimulated | 121.4 |
| MyLa (CD8+ T cells) | 130.0 |
| Naïve T cell | 151.9 |

**Figure S1.**
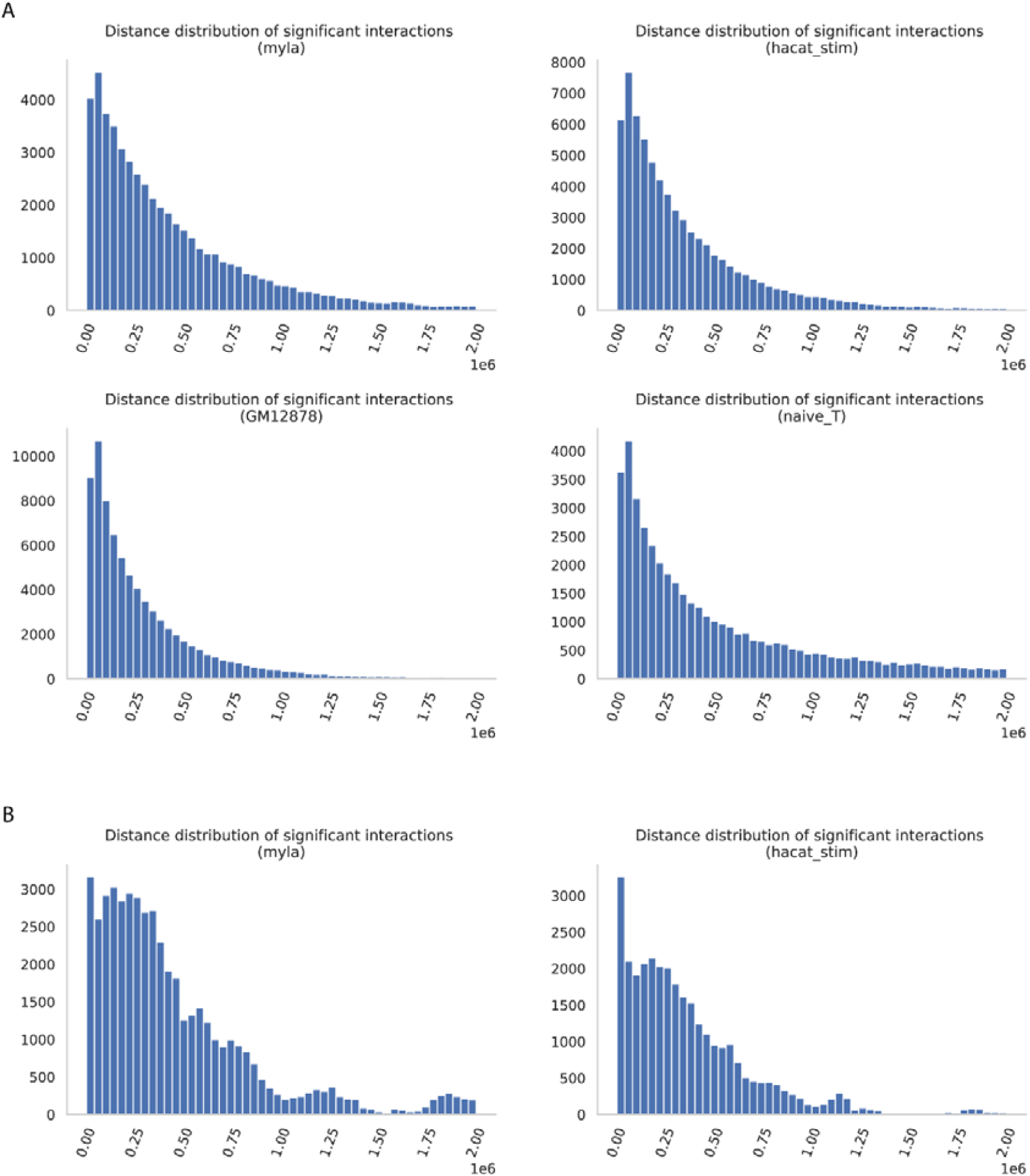
Distribution plots of significant interactions. A) HiChIP; B) Capture-Hi-C.

**Figure S2.**
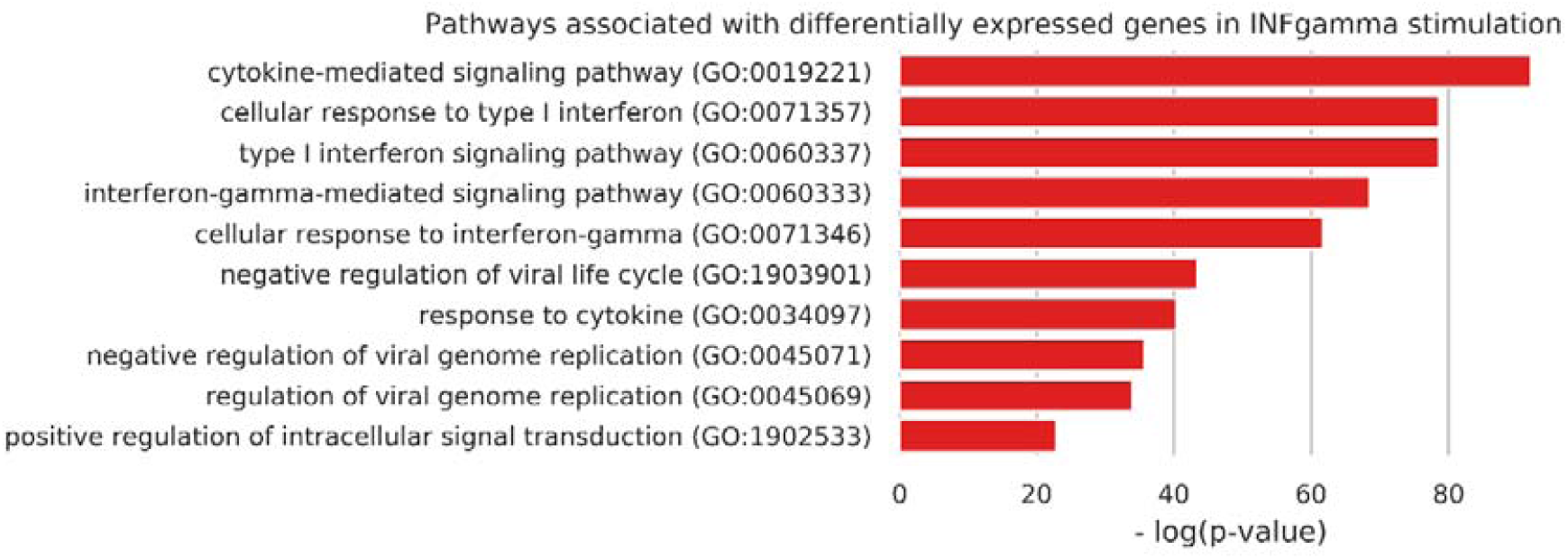
Pathways identified by DE genes it IFN-γ stim.

**Figure S3.**
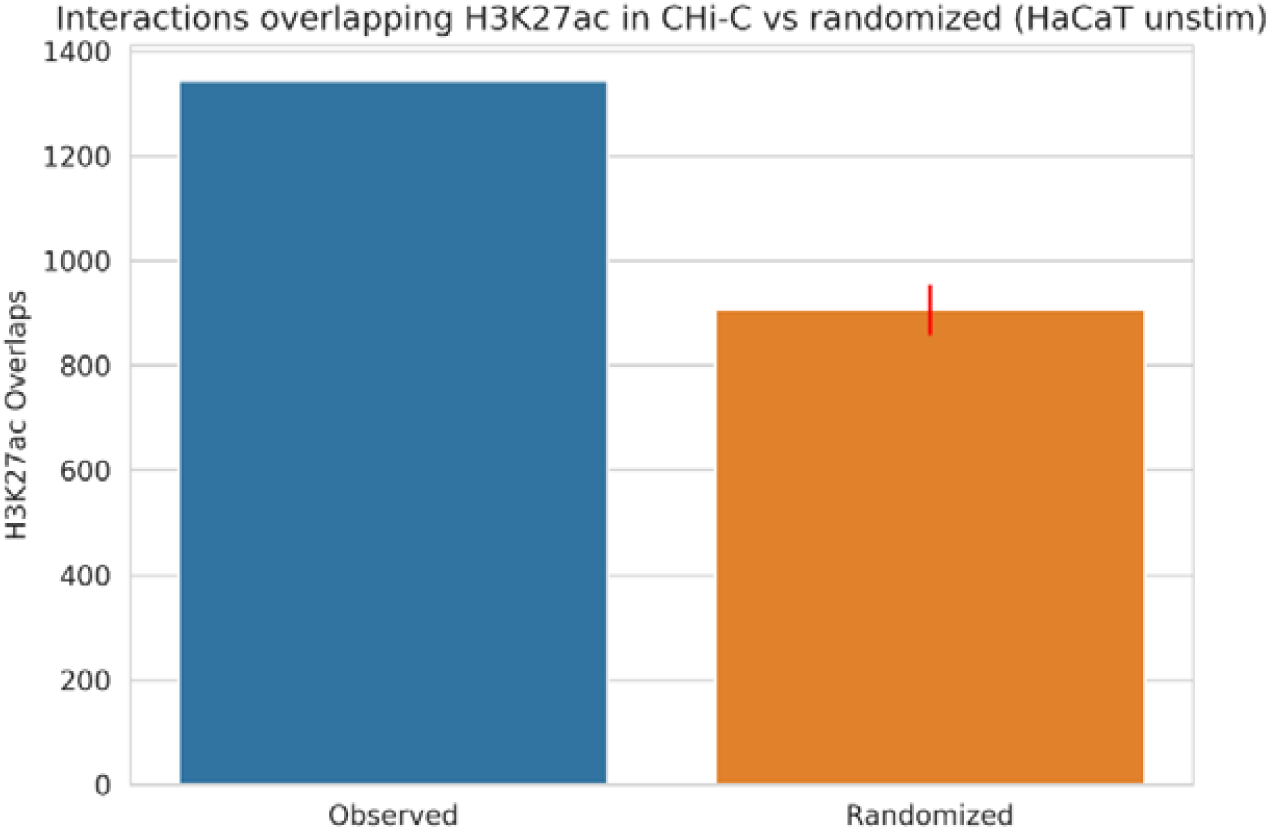
Enrichment of H3K27ac peaks in significant Capture-Hi-C interactions vs randomized. Red line indicates 95% confidence interval.

**Figure S4.**
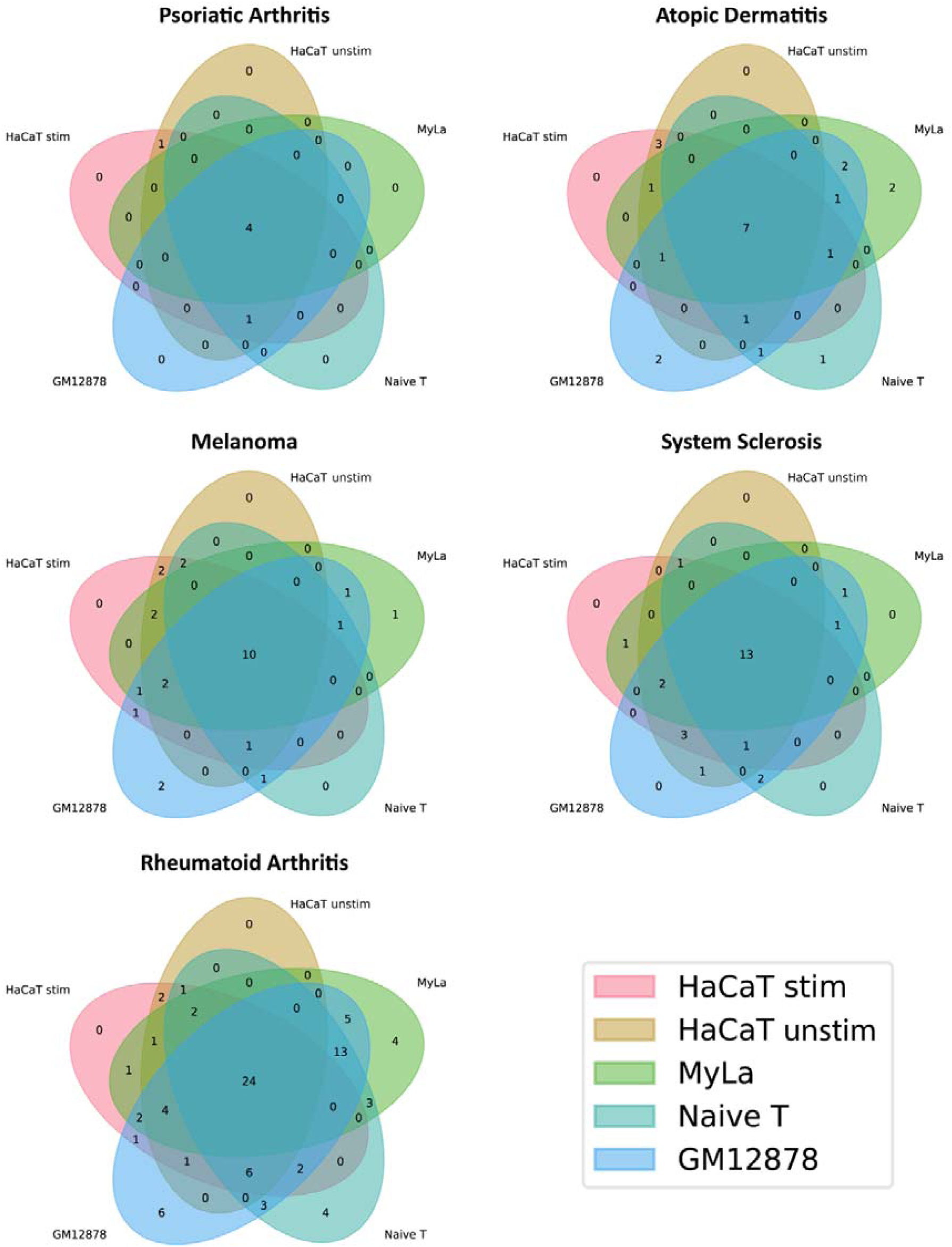
Venn diagram showing the number of loci which have at least one variant overlapping a HiChIP H3K27ac peak in each cell type for other diseases.

**Figure S5.**
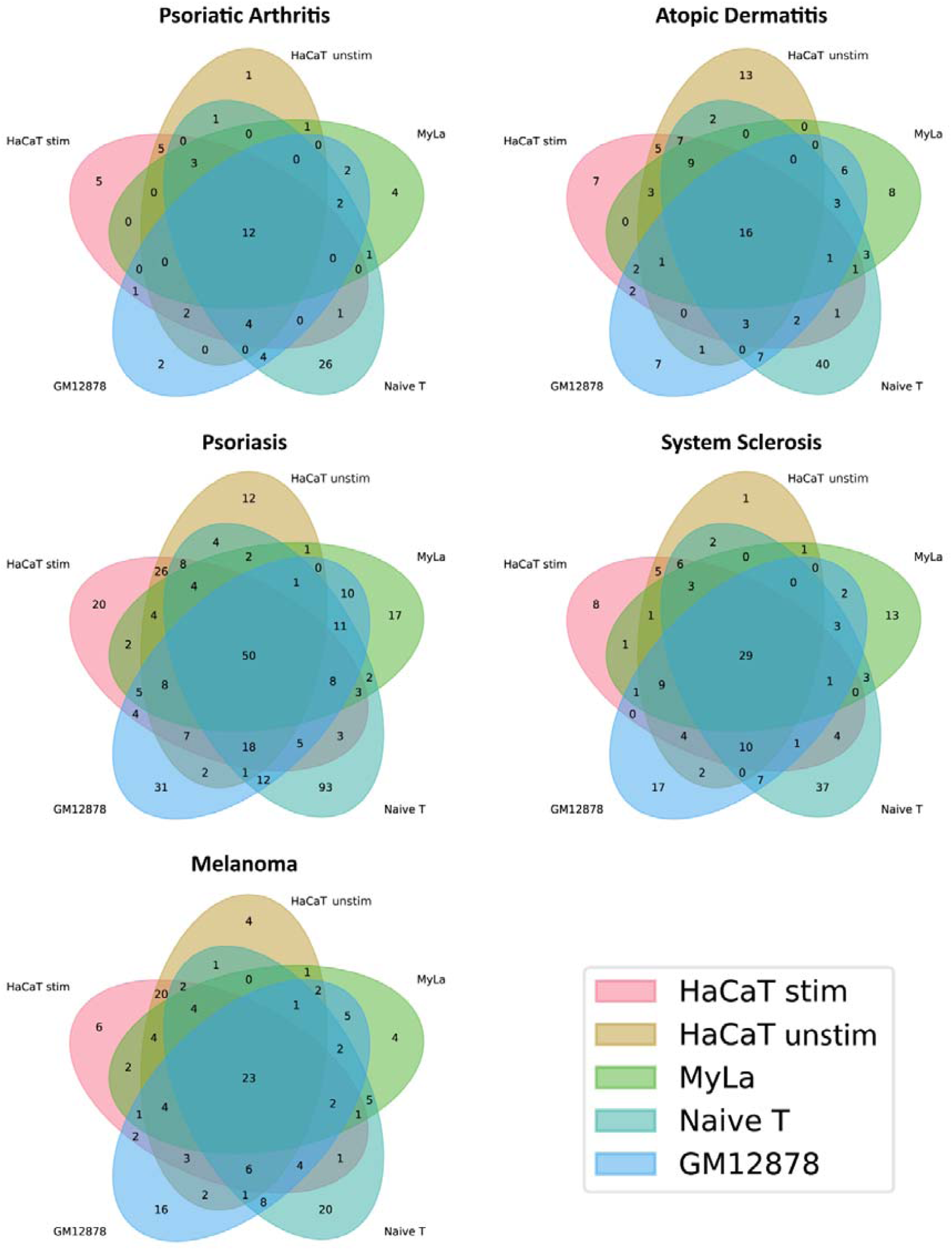
Venn diagram representing the number of genes identified by cell type.

**Figure S6.**
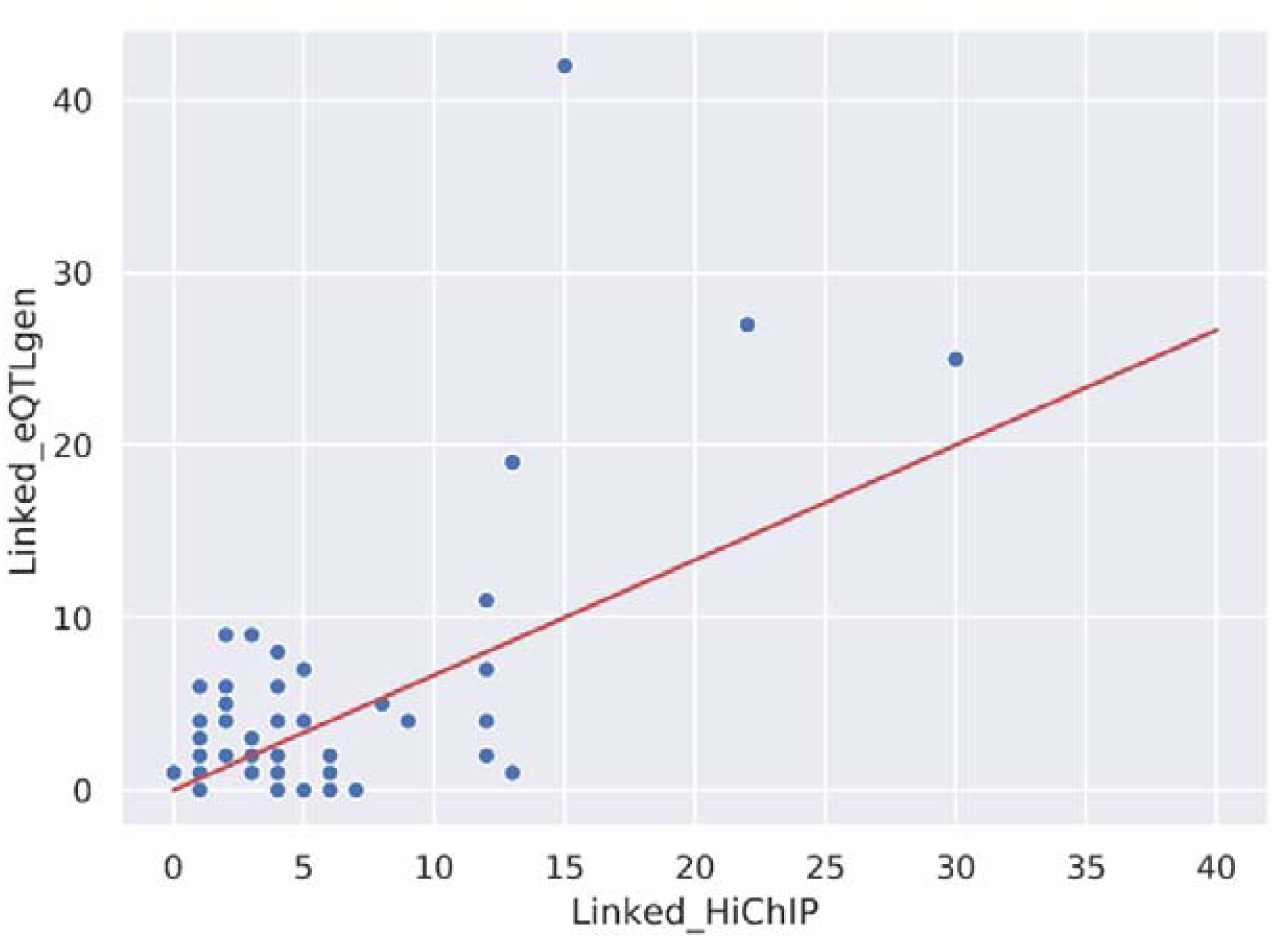
The number of genes linked by HiChIP in Naïve T cells and GM12878 cells correlates with the number of genes linked by eQTL from eQTLgen (R^2^ = 0.631,).

**Figure S7.**
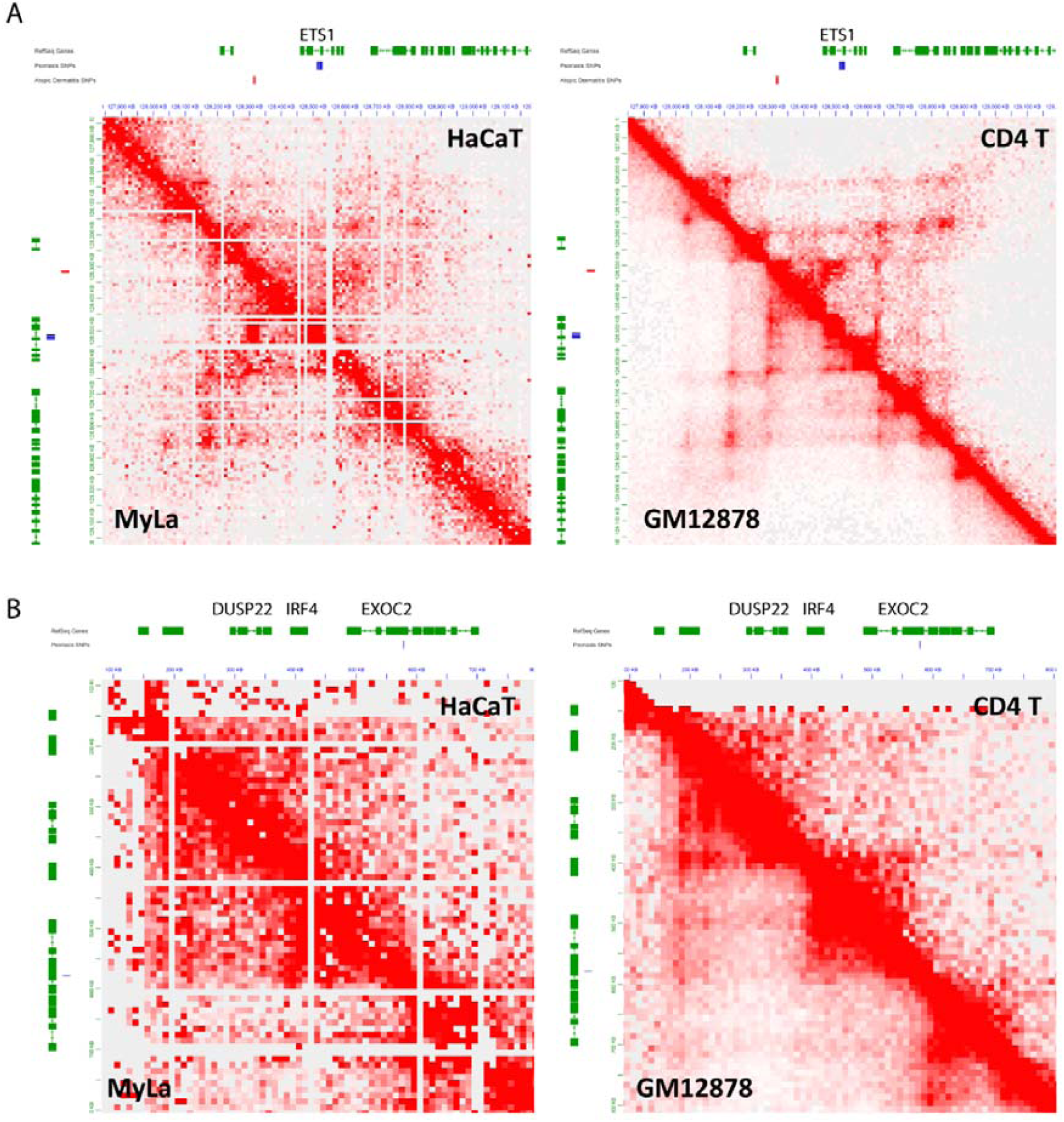
A) Hi-C contact maps for the ETS1 locus; B) Hi-C contact maps for the EXOC2/IRF4/DUSP22 locus.

**Figure S8.**
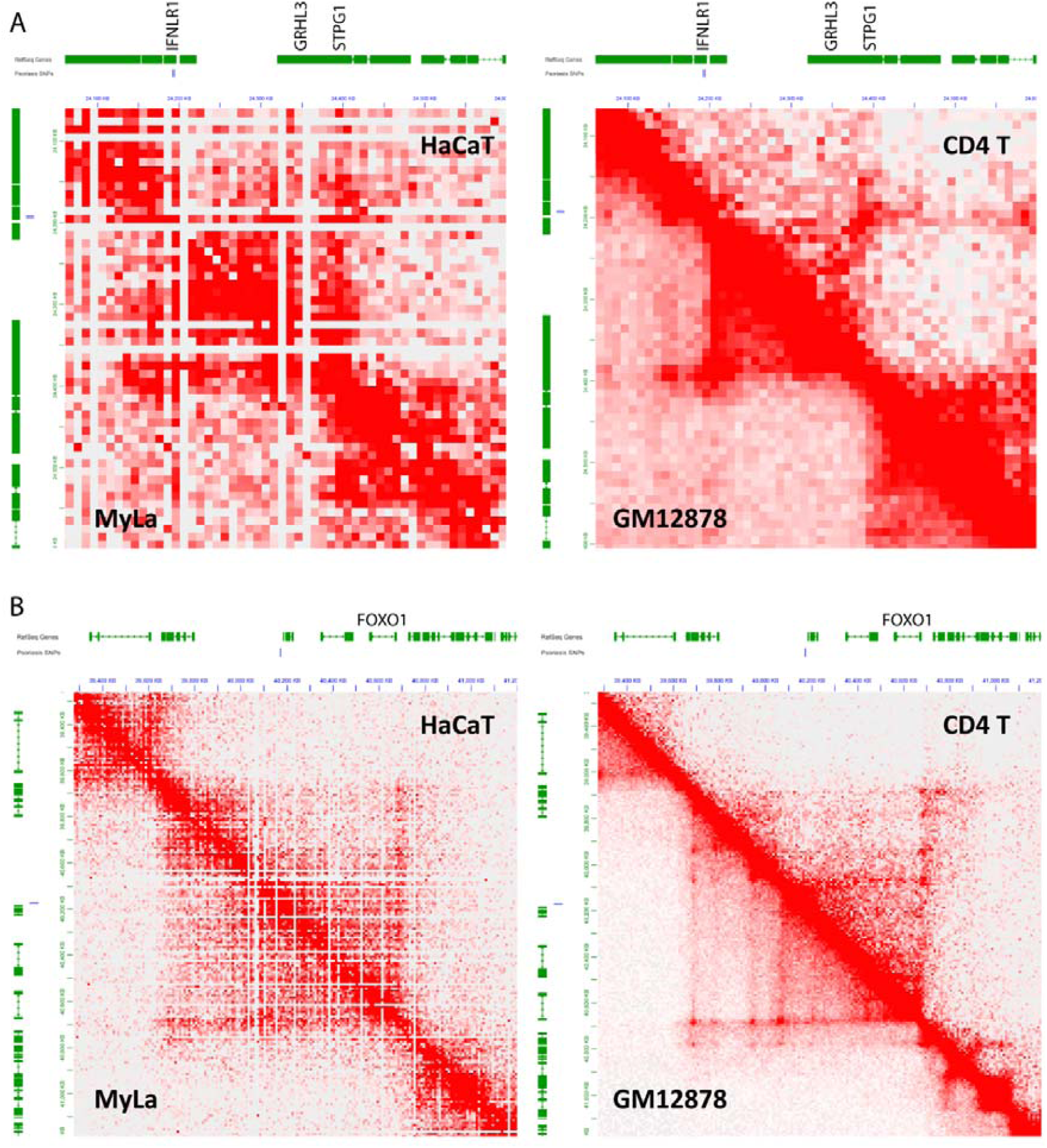
A) Hi-C contact maps for the IFNLR1/GRHL3 locus; B) Hi-C contact maps for the FOXO1 locus.

**Figure S9.**
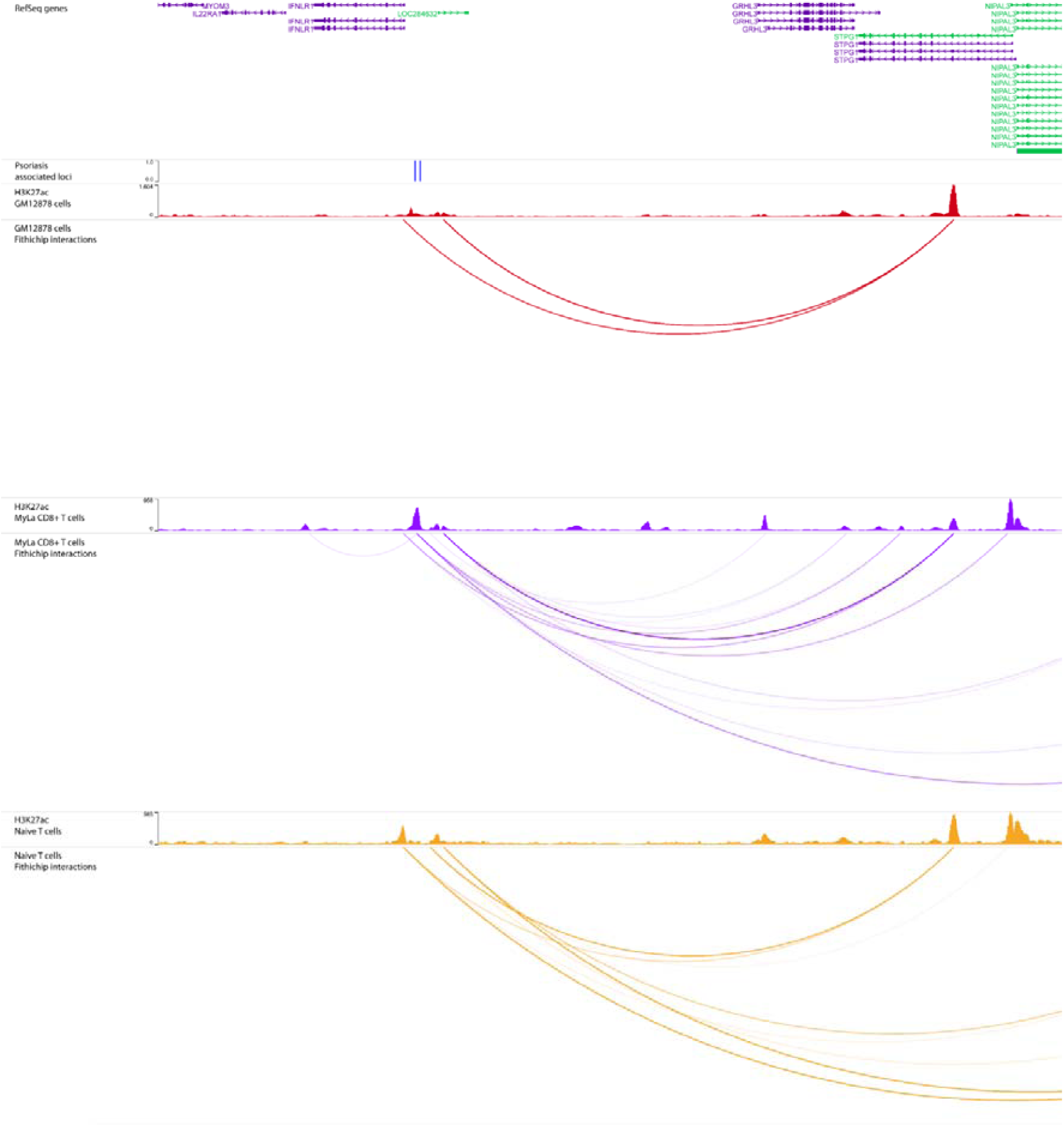
HiChIP interactions from the IFNLR1/GRHL3 locus link GRHL3 as a candidate gene in psoriasis. Tracks for other cell types. Tracks (in order): RefSeq genes; SNPs associated with Psoriasis; H3K27ac signal in GM12878 cells; Significant long range interactions originating from the psoriasis associated locus in GM12878 cells; H3K27ac signal in MyLa cells; Significant long range interactions originating from the psoriasis associated locus in MyLa cells; H3K27ac signal in Naïve T cells; Significant long range interactions originating from the psoriasis associated locus in Naïve T cells.

